# KEAP1 mutation in lung adenocarcinoma promotes immune evasion and immunotherapy resistance

**DOI:** 10.1101/2021.09.24.461709

**Authors:** Anastasia-Maria Zavitsanou, Yuan Hao, Warren L. Wu, Eric Bartnicki, Ray Pillai, Triantafyllia Karakousi, Alberto Herrera, Angeliki Karatza, Ali Rashidfarrokhi, Sabrina Solis, Metamia Ciampricotti, Anna H. Yeaton, Ellie Ivanova, Corrin A. Wohlhieter, Terkild B. Buus, John T. Poirier, Charles M. Rudin, Kwok-Kin Wong, Andre L. Moreira, Kamal M. Khanna, Aristotelis Tsirigos, Thales Papagiannakopoulos, Sergei B. Koralov

## Abstract

Lung cancer treatment has benefited greatly from the development of effective immune-based therapies. However, these strategies still fail in a large subset of patients. Tumor-intrinsic mutations can drive immune evasion via recruiting immunosuppressive populations or suppressing anti-tumor immune responses. *KEAP1* is one of the most frequently mutated genes in lung adenocarcinoma patients and is associated with poor prognosis and inferior response to all therapies, including checkpoint blockade. Here, we established a novel antigenic lung cancer model and showed that *Keap1*-mutant tumors promote dramatic remodeling of the tumor immune microenvironment. Combining single-cell technology and depletion studies, we demonstrate that *Keap1*-mutant tumors diminish dendritic cell and T cell responses driving immunotherapy resistance. Importantly, analysis of *KEAP1* mutant patient tumors revealed analogous decrease in dendritic cell and T cell infiltration. Our study provides new insight into the role of *KEAP1* mutations in promoting immune evasion and suggests a path to novel immune-based therapeutic strategies for *KEAP1* mutant lung cancer.

**Statement of significance:** This study establishes that tumor-intrinsic *KEAP1* mutations contribute to immune evasion through suppression of dendritic cell and T cell responses, explaining the observed resistance to immunotherapy of *KEAP1* mutant tumors. These results highlight the importance of stratifying patients based on *KEAP1* status and paves the way for novel therapeutic strategies.

## Introduction

Lung cancer is the leading cause of cancer-related deaths worldwide(1). Non-small cell lung cancer (NSCLC), including lung adenocarcinoma (LUAD), accounts for 85% of lung cancer cases(2,3). Despite improvements in therapy, NSCLC mortality remains high with fewer than 20% of patients surviving after 5 years(2). In recent years immunotherapy has emerged as an important therapeutic intervention for a wide range of cancer types, including NSCLC (4). Immunotherapeutic approaches include the use of monoclonal antibodies against major checkpoints, such as cytotoxic T-lymphocyte antigen (CTLA4) and programmed death receptor 1 (PD1) or its ligand PDL1 (reviewed in (5)). Even though checkpoint inhibitors, either alone or in combination with chemotherapy, have shown great promise in the context of NSCLC, improvements in progression free and overall survival are relatively modest, with only a subset of patients exhibiting long lasting remissions (5-8). Understanding what limits the efficacy of these treatments is critical for development of future therapeutic approaches.

Emerging evidence suggests that a major factor that can impact immunotherapy response is whether the tumor displays an immunologically “cold” or “hot” tumor immune microenvironment (TIME) (9). Cytotoxic T lymphocytes (CTLs) are main drivers of the anti-tumor immune response and their presence is associated with improved outcome and response to checkpoint blockade therapies (10,11). Accordingly, CTL chemoattractants such as CXCL9/CXCL10 and cytokines like type I and type II interferons have also been associated with favorable outcome, as they can promote CTL effector function (12-14). In addition, Batf3-dependent conventional type I dendritic cells (cDC1) can modulate anti-tumor T cell responses via chemokine production and presentation of tumor-associated antigens, together with providing co-stimulatory or inhibitory signals (15-17). Importantly, cDC1 have been shown to control checkpoint responses in preclinical models and correlate with improved survival in humans (18-21). The full spectrum of tumor-intrinsic and host-dependent factors that determine the strength of anti-tumor immune response remain to be fully elucidated.

One approach to address the challenge of immunotherapy resistance is to define tumor-intrinsic genetic mutations that can modulate the TIME and regulate response to therapy. Oncogenic pathways can modulate anti-tumor immune responses via promoting the recruitment of immunosuppressive populations such as neutrophils (22) or suppressing antigen-specific T cell responses (13), thereby establishing an immunologically cold TIME.

After *KRAS* and *TP53, KEAP1* (Kelch-like ECH-associated protein 1) is the third most frequently mutated gene in LUAD (23). KEAP1 is a protein adaptor providing specificity for the CUL3 (Culin 3)/RBX1 (Ring box 1) E3 ubiquitin ligase complex. KEAP1 negatively regulates the antioxidant transcription factor NRF2, constitutively targeting it for proteasomal degradation, thereby preventing nuclear accumulation and activation of the antioxidant program (24-26). The majority of somatic mutations in *KEAP1* are missense or truncating events that can generate dominant negative forms of *KEAP1* (27-30). We previously established that *KEAP1* is a tumor suppressor gene by demonstrating that *Keap1* loss accelerated *Kras*-driven LUAD progression (31) and metastasis (32). Emerging data indicates that *KEAP1* mutant lung cancers are resistant to checkpoint inhibition (33,34) and associate with a “cold”, lacking T cells tumor microenvironment (35). This suggests that in addition to tumor-intrinsic effects, mutations in *KEAP1* impact cancer immune surveillance.

In this study, we investigated whether tumor-intrinsic *Keap1* mutations remodel the TIME and promote resistance to checkpoint blockade therapies. We established a novel H-Y driven antigenic lung cancer model that is able to promote adaptive T cell immunity and responds to anti-PD1 therapy. We found that *Keap1* mutation drives downregulation of interferon gene signatures and expression of chemokines critical for anti-tumor immunity. Using both single-cell technology and depletion studies, we demonstrate that *Keap1*-mutant tumors suppress cDC1 mediated CD8 T cell immunity and drive resistance to immunotherapy. To assess the translational relevance of our findings, we examined the immune microenvironment of *KEAP1* wild-type and mutant human LUAD tumors using multi-color immunofluorescence and high-plex *in situ* proteomic analysis. We showed that *KEAP1*-mutant human tumors display decreased T cell and DC infiltration and negatively correlate with genes associated with checkpoint blockade responses. Taken together, our study reveals how tumor-intrinsic *KEAP1* mutations subvert anti-tumor immune responses and underscores the importance of stratifying LUAD patients based on *KEAP1* mutation status prior to selection of immunotherapy regimen.

## Results

### *Keap1* mutation accelerates tumor growth in antigenic and autochthonous mouse models of lung adenocarcinoma

Given that previous studies have implicated tumor-intrinsic *KEAP1* mutations in failure to respond to immune checkpoint inhibitors, we sought to examine the impact of *KEAP1* mutations on cancer growth and immune surveillance. We generated an orthotopic transplant model of Kras^G12D/+^; p53^-/-^ (KP) lung cancer (36) with cells carrying either a wild-type or a dominant negative form of *Keap1* (Keap1 R470C), the most frequently listed mutation of *KEAP1* in the COSMIC database (Fig.1A; Fig. S1A) (29,37). Because earlier work has demonstrated that Kras-driven mouse LUAD models do not elicit robust T cell responses and do not respond to checkpoint inhibition therapy (38-42), we elected to establish a Y chromosome-driven antigenic mouse model of LUAD. We utilized gene-targeted male tumor cells (Fig. S1B) and introduced them orthotopically into immunocompetent female hosts (Fig 1A). We hypothesized that the presence of Y chromosome antigens will elicit an antigenic response in female recipients as previously shown in the context of lymphoma and liver cancer (43,44). We validated our H-Y-driven system by ensuring that male KP lung adenocarcinoma cells trigger an anti-tumor immune response in females but not in males. Indeed, KP cells grew rapidly in male mice, while tumor growth was controlled in female hosts (Fig.1B). We then performed antibody-mediated depletion of CD4 or CD8 T cells in female and male hosts. Depletion of either CD4 or CD8 T cells resulted in increased tumor burden in female (Fig.1C), but not in male mice (Fig.1D).

**Figure 1:**
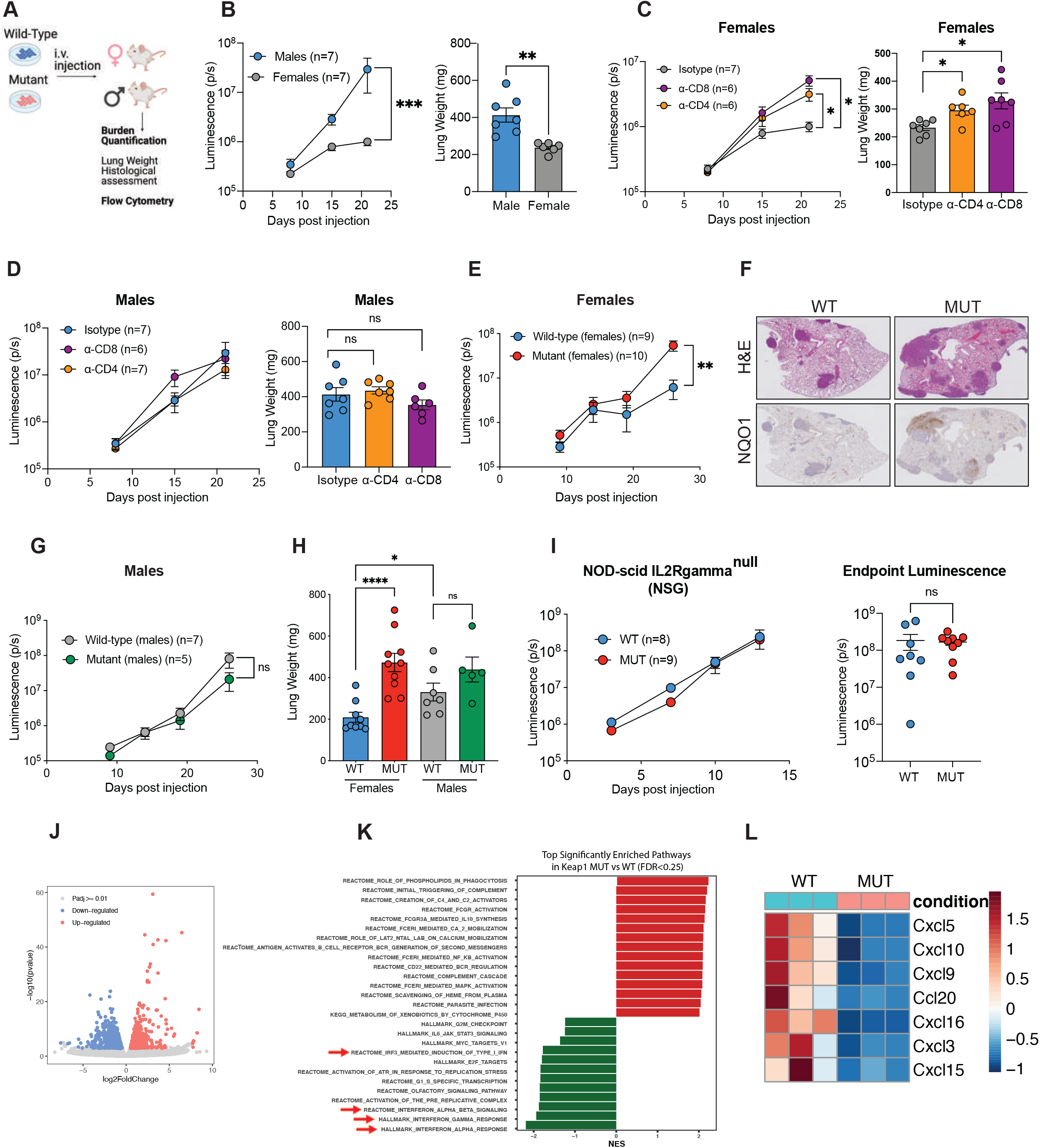
Loss of *Keap1* promotes immune evasion and accelerated tumor growth in a novel antigenic model of LUAD. (A) Schematic representation of our antigenic H-Y-driven orthotopic mouse model. (B) Growth kinetics of male KP tumors established in female or male hosts. (C) Growth kinetics (Left) of male KP cells grown in female hosts following antibody-mediated depletion of CD4+ or CD8+ T lymphocytes. Right: Lung weight measurement of whole lungs as a proxy for tumor burden (104) (D) Growth kinetics (Left) of male KP cells injected into male hosts following antibody mediated depletion of CD4+ or CD8+ T lymphocytes. Right: Lung weight measurements of whole lungs. (E) Growth kinetics of *Keap1* wild-type and mutant KP cells in females. (F) Representative images of H&E and NQO1 immunohistochemical staining of *Keap1* wild-type and mutant KP tumors in female hosts. (G) Growth kinetics of *Keap1* wild-type and mutant KP cells injected in male hosts. (H) Lung weight as proxy for tumor burden measured on day 26 in female and male mice bearing *Keap1* wild-type and mutant tumors (nζ5 mice per group). (I) Growth kinetics (left) and endpoint luminescence (right) of *Keap1* wild-type and mutant male cells injected into immunodeficient (NSG) mice. (J) Volcano plot showing differentially expressed genes between wild-type and mutant tumor cells. (K) Most significantly enriched or depleted pathways (FDR <0.25) in *Keap1*-mutant compared to wild-type tumor cells isolated via sorting from tumor-bearing female hosts and subjected to RNA-seq. (L) Gene expression heatmap of chemokines significantly (p<0.05) altered between *Keap1*-mutant and wild-type tumor cells. Growth kinetics were monitored using *in vivo* luminescence at specified timepoints. *P<0.05; **P<0.01; ***P<0.001; ****P < 0.0001

To investigate the role of *Keap1* mutation in modulating anti-tumor immune responses, we injected *Keap1* wild-type and mutant male mouse lung cancer cells into immunocompetent female hosts (Fig. 1E). Critically, *Keap1-*mutant tumors displayed accelerated growth kinetics compared to wild-type tumors in female mice (Fig. 1E), but no difference in tumor growth based on *Keap1* mutation was observed in male recipients (Fig. 1G,H; FigS1F). Growth differences between wild-type and mutant tumors were observed after day 15 suggesting involvement of adaptive immune responses (Fig. 1E). Histological analysis showed increased tumor burden in the lungs of female mice injected with *Keap1*-mutant cells (Fig. 1F). To validate loss of KEAP1 function in our dominant negative model we conducted immunohistochemical analysis for NQO1, as it has been previously shown that *Keap1* loss results in activation of the NRF2 pathway (31,45,46). Our staining confirmed that *Keap1*-mutant tumors expressed higher levels of NQO1, a downstream target of NRF2 (Fig. 1F).

Assessment of *in vitro* proliferative capacity of *Keap1* wild-type and mutant cells did not indicate any differences, suggesting that *Keap1* mutation does not impact the intrinsic proliferative capacity of lung adenocarcinoma cells (Fig. S1C). To control for *Keap1* overexpression, we also generated lung adenocarcinoma cells carrying an empty control vector. The *in vitro* and *in vivo* growth of tumor cells transduced with an empty vector was similar to cells transduced with *Keap1* wild-type vector (Fig. S1B-E). Together, these findings suggest that *Keap1* mutations elicit faster tumor growth by altering the host immune response.

If *Keap1* mutation promotes accelerated tumor growth via suppression of anti-tumor immune responses, then *Keap1*-mutant tumors should grow similarly to *Keap1* wild-type tumors in a non-antigenic setting or in immunocompromised mice. Indeed, we did not observe any differences in growth of wild-type and mutant cells when injected into male hosts (Fig.1G,H;FigS1F) or in NOD-SCID IL2Rgamma^null^ (NSG) mice that lack B, T, and NK cells (Fig. 1I).

We next sought to investigate how *Keap1* mutation impacts tumor growth and immune surveillance in an autochthonous genetically engineered mouse model (GEMM) of LUAD. We generated Kras LSL-^G12D/+^; Trp53^flox/flox^ (KP) mice expressing either wild-type *Keap1* (*Keap1*^*+/+*^), mice heterozygous for the floxed *Keap1* conditional allele (*Keap1*^*fl/+*^) or animals homozygous for the conditional allele (*Keap1*^*fl/fl*^) (Fig. S1G). Intratracheal administration of *Cre*-expressing lentivirus leads to the induction of LUAD tumors in these KP animals. Tumor burden quantification based on total lung weight and histological analysis revealed that partial loss of *Keap1* resulted in increased tumor burden (Fig. S1H-J). Interestingly, mice in which *Cre-*mediated deletion resulted in homozygous loss of *Keap1* had similar tumor burden to animals with a WT *Keap1* allele (Fig. S1H-J), which is consistent with prior studies (46) and data from human tumors suggesting that more aggressive tumors first acquire a missense or truncating mutations in one allele and then undergo loss of heterozygosity (27,28). Immunohistochemical staining for NQO1 displayed the expected gradient, where *Keap1*^*+/+*^ tumors had little to no staining, *Keap1* ^fl/+^ tumors had intermediate staining, and *Keap1*^fl/fl^ were strongly positive (Fig. S1H, K).

### Keap1 mutation results in down regulation of genes critical for anti-tumor immunity

To investigate the molecular mechanisms by which *Keap1*-mutant lung tumors could be driving immune evasion, we sorted tumor cells from the lungs of immunocompetent female mice with KP tumors carrying a wild-type or *Keap1*-mutant allele. We then performed unbiased transcriptional profiling of the sorted tumor cells followed by differential gene expression analysis. We found that 655 genes were upregulated while 1108 were downregulated in mutant compared to wild-type cells (Fig.1J). Pathway enrichment analysis of our RNA sequencing (RNA-seq) data revealed that interferon pathways were the top down-regulated pathways in *Keap1*-mutant compared to wild-type tumor cells (Fig 1K). Interferon production has been shown to be critical for activation of cross presentation of tumor antigens by DCs to CD8 T cells, as well as for T cell recruitment into the tumor microenvironment (15,47,48).

Immune cell infiltration is a key parameter in cancer prognosis and response to immune therapies. Dysregulated chemokine signaling may contribute to immune evasion, as chemokines are essential for migration of both activating and immunosuppressive leukocytes in the TIME (12-14,16,22,49-52). We examined chemokine expression in wild-type and mutant tumor cells. A total of seven chemokines were differentially expressed (Cxcl9, Cxcl10, Cxcl16, Ccl20, Cxcl3, Cxcl5, Cxcl15) with all of them being expressed at lower levels in *Keap1*-mutant compared to wild-type tumor cells (Fig. 1L). The reduced expression of these chemokines can contribute to exclusion of host immune responses including decreased infiltration of key anti-tumor effector cells.

### Keap1 mutation prevents CD103 DC (cDC1) accumulation and activation in lung adenocarcinoma tumors

We hypothesized that decreased interferon signaling and chemokine production could result in extensive remodeling of TIME in *Keap1*-mutant tumors. To investigate how Keap1 mutation alters the immune microenvironment, we performed single-cell RNA-seq (scRNA-seq) of wild-type and mutant lung tumors. We used flow cytometric sorting to enrich for lung tissue-infiltrating immune cells, being sure to exclude circulating leukocytes using intravenous injection of fluorescently-tagged CD45 antibody. We identified 14 discrete clusters in our scRNA-seq analysis representing all major immune cell lineages with granulocytes, T lymphocytes, B cells, macrophages and dendritic cell subsets being the most abundant populations (Fig. S2A-D). Given the critical role DCs and T cells in anti-tumor immunity, we wanted to investigate how *Keap1* mutation is impacting DC mediated T cell immunosurveillance in LUAD.

Tissue resident dendritic cells consist of two subsets: Batf3-dependent CD103+/CD8+ DCs (cDC1) specialized in cross presenting antigens to CD8 T cells through class I major histocompatibility complex (MHC) pathway (53) and CD11b+ DCs (cDC2) which drive CD4+ helper T cell responses (54,55). cDC1 have been shown to be essential for anti-tumor immune responses in multiple preclinical mouse models and were recently shown to promote anti-cancer immunity in the KP lung cancer model (15-19,41,47,48,56).

To get a more detailed look at TIME-associated DC populations, we performed dendritic cell sub clustering of our scRNA-seq dataset. We identified 6 different clusters: CD103 DCs, proliferating CD103 DCs, CD11b DCs, proliferating CD11b DCs, plasmacytoid dendritic cells and DCs with migratory features, which seem to correspond to the most recently characterized mreg DCs (Fig. 2A) (41). CD103 DCs cluster expressed canonical genes *Itgae, Xcr1, Clec9a, Naaa*, while CD11b DC characteristic transcripts included *Itgam, Mgl2, CD209a (57)*. In addition to bona fide cDC1 and cDC2 genes, the proliferating cDC1 and cDC2 clusters also expressed cell cycle and proliferation markers *Birc5, Mki67, Top2a, Stmn1, Cks1b, Hist1h2ae, Hist1h1b, Hist1h2ap*. The mreg DC cluster expressed *Ccr7, Fscn1, Il4i1* as previously shown (41). pDCs expressed canonical genes *Bst2, Siglech, Ccr9, Ly6d, Klk1* (Fig. S2E) (58). DC subclustering revealed that *Keap1*-mutant tumors are characterized by decreased presence of CD103 DCs, increased frequency of pDCs and modestly increased CD11b and mreg-DC numbers (Fig. 2B, C).

**Figure 2:**
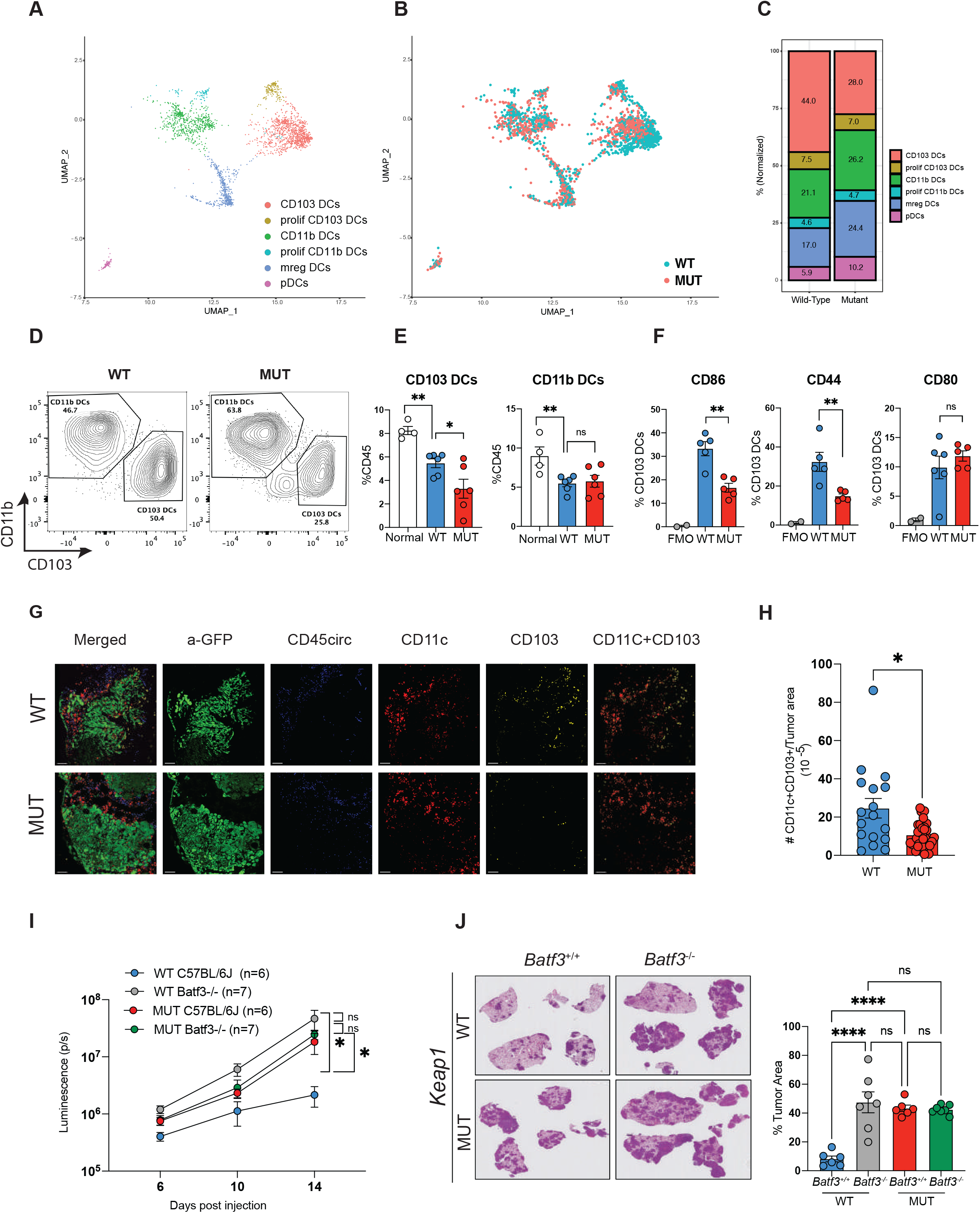
Absence of cDC1 mediated anti-tumor immune responses in *Keap1*-mutant tumors. (A) UMAP visualization of DC sub-clusters identified by scRNA-seq, clustered and colored by cell type. Clusters identified based on gene expression. (B) UMAP representation of the distribution of DC subclusters in *Keap1* wild-type (blue) and mutant (red) lung tumors. (C) Bar plot showing distribution of the different DC clusters in *Keap1* wild-type and mutant mouse lung tumors. (D) Representative flow cytometry plots for CD103 versus CD11b within a CD11c+MHCII+ DC gate. (E) Percentage of CD103 and CD11b DCs out of total tissue-infiltrating immune cells (CD45+CD45circ-) in normal (non-tumor) lung and lungs with *Keap1* wild-type and mutant tumors. Each symbol represents an individual mouse. (F) Percentage of CD86, CD44 and CD80 among CD103 DCs in wild-type and mutant *Keap1* tumors. FMO (fluorescence-minus-one) control shown for all markers. Each symbol represents an individual mouse (G) Confocal images of lung tumor sections from *Keap1* wild-type and mutant KP tumors. GFP signal from tumors is shown in green, circulating CD45+ cells shown in blue, CD11c shown in red and CD103 shown in yellow. Scale bars shown are 100 μm. Images are representative of individual tumors from ≥ 5 mice per genotype. (H) Quantification of tissue infiltrating CD103 DCs (CD11c+CD103+CD45circ-) in tumor (GFP+) areas. Each symbol represents an individual tumor. (I) Growth kinetics of *Keap1* wild-type and mutant lung orthotopic tumors in C57BL6J and *Batf3*^-/-^ female mice. Tumor cells express luciferase enabling monitoring of tumor growth kinetics via luminescence. (J) Quantification of tumor burden (tumor area/total lung area) of *Keap1* wild-type and mutant tumors in C57BL/6J and Batf3-/-based on H&E staining. Each experimental subgroup had at least n=6 mice. *P<0.05; **P<0.01; ***P<0.001; ****P < 0.0001

We optimized flow cytometric staining strategy that allows distinction between cDC1 and other myeloid immune populations, including macrophages and cDC2 (Fig.S3A) and confirmed that CD103 DCs were decreased in *Keap1*-mutant compared to wild-type tumors while CD11b+ DCs remained unchanged (Fig. 2D, E). Importantly, cDC1 infiltration was lower in *Keap1*^fl/+^ compared to *Keap1*^+/+^ autochthonous model (Fig.S3B) indicating that *Keap1* mutation suppresses cDC1 infiltration in both orthotopic and autochthonous lung cancer models. Consistent with the immunosuppressive role of *Keap1* mutation, there was no difference in cDC1 infiltration between wild-type and mutant tumors in the less antigenic setting of male hosts (Fig. S3C). Finally, when we examined markers of DC activation, we observed that cDC1 from *Keap1* wild-type tumors displayed higher levels of co-stimulatory molecules such as CD44 and CD86 compared to dendritic cells from mutant tumors (Fig. 2F).

To examine the distribution of cDC1 in *Keap1* wild-type and mutant tumors, we conducted multi-color immunofluorescence staining. We observed that cDC1 were primarily found in tumor periphery (Fig. 2G). Interestingly, *Keap1* wild-type tumors presented increased intratumoral infiltration (Fig. 2G) and higher overall cDC1 numbers (Fig. 2H) compared to *Keap1*-mutant tumors.

We reasoned that if *Keap1*-mutant tumors suppress cDC1 mediated anti-tumor immune responses, then *Keap1* wild-type and mutant tumors should grow similarly in cDC1-deficient Batf3^-/-^ hosts. Using Batf3^+/+^ and Batf3^-/-^ female C57BL/6J hosts we showed that, although mutant tumors displayed accelerated growth in WT mice, the difference in growth kinetics between these tumor genotypes is not evident in Batf3^-/-^ hosts (Fig. 2I-J). Based on histological assessment, cDC1 deficiency resulted in increased tumor burden in mice with KP tumors while *Keap1-*mutant tumors displayed similar growth kinetics in the presence or absence of cDC1 subset (Fig. 2K). These results suggest that *Keap1*-mutant tumors suppress cDC1 mediated immune surveillance and highlight the central role of cDC1 population in the coordination of anti-tumor immune responses.

### Keap1 mutation suppresses cDC1-mediated CD8 T cell immunity

cDC1 are responsible for taking up dead tumor cells and cross priming CD8 and CD4 T lymphocytes(18,53,59,60). In addition to this role they can also promote recruitment and local expansion of CD8 T lymphocytes and support their effector function (61). We thus sought to determine whether reduced number and functionality of cDC1 observed in *Keap1*-mutant tumors is driving defective T cell responses.

Upon sub clustering of T lymphocytes in our scRNA-seq dataset we identified 10 phenotypically distinct populations of T cells based on gene expression profiles(Fig. 3A). Most of these clusters represented cells at different stages of activation and exhaustion. Based on patterns of gene expression we identified CD8 and CD4 naïve cells (Sell^+^Ccr7^+^Dapl1^+^Igfbp4^+^), early activated CD8 and CD4 T cells characterized by intermediate expression of *Sell, Ccr7, Lef1* and upregulation of activation markers common to both CD4 and CD8 T cells (*Ccl5, Nkg7*), or unique to either CD8 (*Klrk1, Xcl1, Cd7, Ly6c2*) or CD4 T cells (*Cd28*) (Fig.3C). We also observed effector/exhausted CD4 and CD8 T cells. These exhausted lymphocytes did not express markers associated with naïve cells, maintained expression of activation markers, but also robustly expressed markers of exhaustion (*Pdcd1, Tigit, Lag3, Bhlhe40*) (Fig. 3A, C). We also identified a CD8 population that appeared to be a cell state between early activated and exhausted cells as it was negative for naïve markers, expressed markers of activation, but no notable expression of makers characteristic of exhausted cells. There were two additional CD4 clusters that appeared to be T regulatory cells (*Il2ra, Ctla4, Klrg1, Foxp3, Tnfrsf4*) and Th1 cells (*Vim, Ahnak, Id2, Crip1, Lgals1, S100a4*) based on gene expression profiles previously associated with these cell phenotypes (Fig. 3A,C) (62,63).

**Figure 3:**
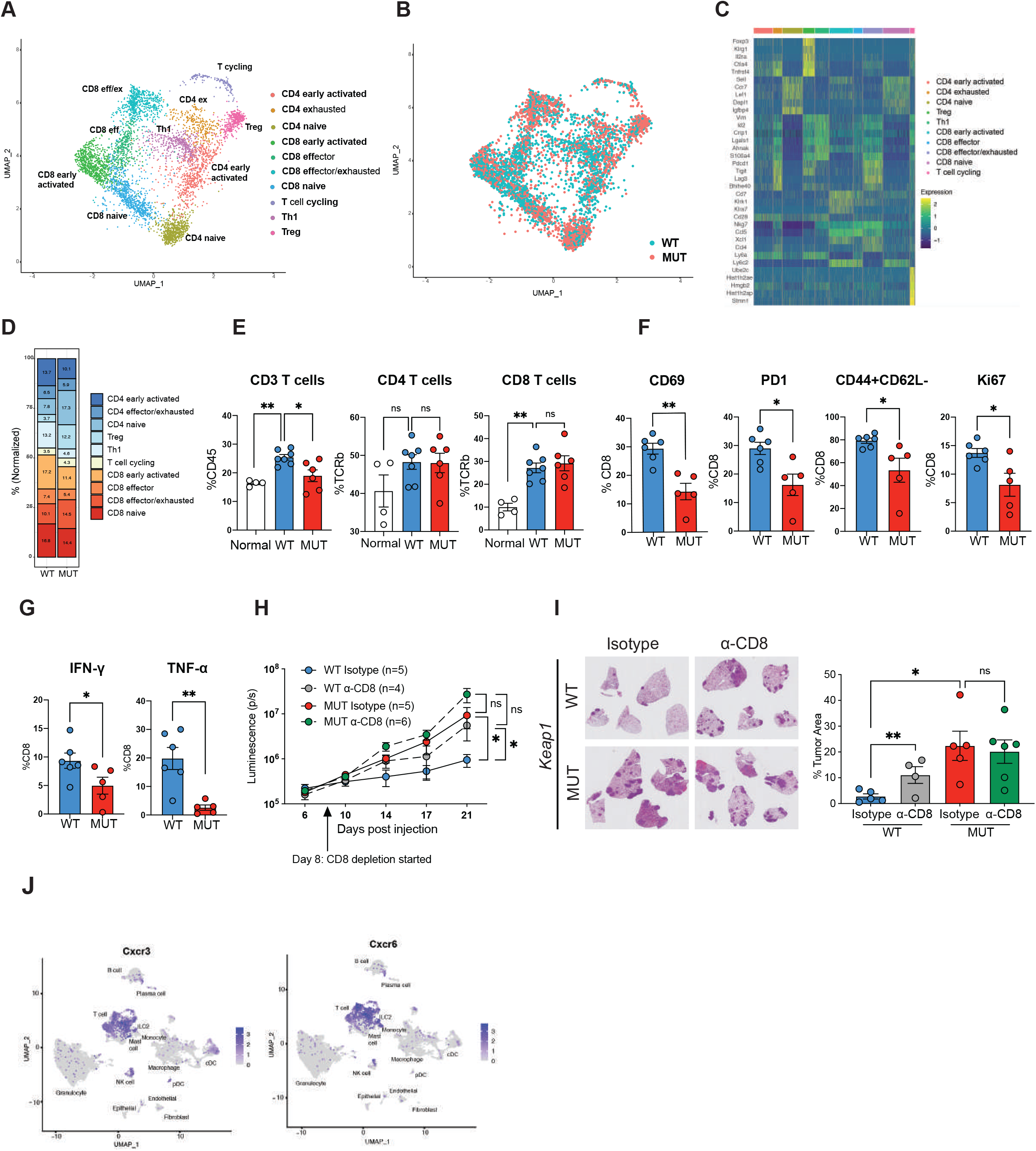
*Keap1*-mutant tumors suppress CD8 T cell responses and promote T cell exhaustion. (A) UMAP visualization of T cell sub-clusters identified by scRNA-seq, clustered and colored by cell type. Clusters identified based on gene expression. (B) UMAP representation of the distribution of T cell subclusters in *Keap1* wild-type (blue) and mutant (red) lung tumors. (C) Heat map shows normalized and log-transferred unique molecular identified (UMI) counts of selected genes, with key indicating cell type of origin. (D) Bar plot showing distribution of the different T cell clusters in *Keap1* wild-type and mutant mouse lung tumors. (E) Percentage of CD3, CD4 and CD8 T cells among tissue-infiltrating immune cells (for CD3) or of tissue-infiltrating TCR β + T cells. Each symbol represents an individual mouse. Each experimental subgroup had n ≥ 4 mice. (F) Percentage of CD69, PD1, CD44+CD62L-, Ki67 among CD8+ T cells in wild-type and mutant *Keap1* tumors. Each symbol represents an individual mouse. Each experimental subgroup had n ≥ 5 mice. (G) Percentage of intracellular IFNγ and TNFα positive cells among the CD8+ T lymphocytes in wild-type and mutant *Keap1* tumors. Each symbol represents an individual mouse. Each experimental subgroup had n ≥ 5 mice. (H) Growth kinetics of *Keap1* wild-type and mutant tumors in female hosts upon antibody-mediated CD8 T cell depletion. Depletion was initiated at Day 8 following verification of tumor engraftment, and continued until experimental endpoint. (I) Representative images of lung tumor burden (left) and quantification (tumor area/total lung area) by H&E staining. Tumor cells express luciferase and thus growth kinetics were monitored using *in vivo* luminescence at specified timepoints *P<0.05; **P<0.01; ***P<0.001; ****P < 0.0001

*Keap1* wild-type tumors had a notable enrichment of both CD4 and CD8 early activated cells (Fig. 3B,D). This is consistent with cDC1 facilitating T cell activation (16,19,20,47,48). On the other hand, *Keap1*-mutant tumors had higher number of CD8 exhausted T cells. Consistent with a more immunosuppressive TME in mutant compared to wild-type tumors, wild-type tumors had increased numbers of Th1 cells while *Keap1*-mutant tumors were enriched in T regulatory cells (Tregs)(Fig 3B,D).

We then performed detailed intracellular and surface flow cytometric analysis (gating strategy Fig. S4A) to distinguish a multitude of T cell subsets in the lungs of tumor-bearing mice. We observed an overall reduction in total T cell infiltration in the lungs of mice with *Keap1-*mutant tumors compared to lungs of animals with KP tumors (Fig. 3E). There was also notable change in the overall landscape of T cells in the lungs of these mice. The difference in number of recruited T cells appears to be driven by a decrease in both CD4 and CD8 T cell subsets in *Keap1-*mutant tumors (Fig. 3E). CD8 T cells infiltrating *Keap1*-mutant tumors expressed lower levels of activation (CD69, PD1, CD44) and proliferation (Ki67) markers (Fig. 3F). Importantly, we observed decreased frequency of CD8 T cells producing the effector cytokines IFN-γ and TNF-α critical for tumor control (Fig. 3G).

When we examined CD4 T cells of wild-type and mutant tumors, we did not observe significant differences in their activation or proliferation, but frequency of cytokine producing CD4 T cells was lower in mutant tumors (Fig. S4B-C). Interestingly, CD4 T cells were skewed towards T regulatory T cells and away from a pro-inflammatory Th1 cell phenotype in *Keap1*-mutant tumors, consistent with the immunosuppressive role of this mutation (Fig. S4D). While there was no difference in total T cells or frequency of Tregs in the less antigenic setting of male hosts (Fig. S4E) we were able to recapitulate the differences in T cells populations noted above in our autochthonous mouse model (Fig. S4F). This suggests that the immunosuppressive phenotype of *Keap1*-mutant tumors is not a consequence of disparity in tumor cell engraftment.

To further investigate whether Keap1 mutation inhibits CD8 and/or CD4 anti-tumor immunity, we performed antibody-mediated depletion of CD8 and CD4 T cells in *Keap1* wild-type and mutant tumors to examine what role Th and cytotoxic T cells play in tumor immune surveillance. Prior to depletion initiation, we confirmed engraftment of tumor cells using *in vivo* bioluminescence imaging to ensure that T cell depletion does not interfere with tumor engraftment. Depletion of CD8 T cells resulted in accelerated growth (Fig. 3H) and increased tumor burden of *Keap1* wild-type tumors without impacting the growth of *Keap1*-mutant tumors (Fig. 3I). This suggests the presence of active CD8 T cell immune surveillance in wild-type, but not *Keap1*-mutant tumors. Absence of CD4 T cells resulted in accelerated growth of both wild-type and mutant tumors, suggesting that there is active CD4 T cell immune surveillance in both tumor genotypes (Fig. S4G-H).

Considering we observed decreased numbers of both T lymphocytes and cDC1 we wanted to investigate whether these immune cell types expressed receptors engaging the chemokines that are downregulated in *Keap1*-mutant cells. Indeed, it was primarily the T cells and cDC1s that expressed CXCR3 that binds to CXCL9/CXCL10 (Fig.3J). T cells also expressed CXCR6 that engages CXCL16 (Fig.3J). Considering that a number of studies have suggested that chemokines CXCL9/10/16 can impact anti-tumor immunity by regulating recruitment of DC and T cells (12,16,64,65)), it is possible that *Keap1*-mutant tumors promote immune evasion via establishing a tumor microenvironment depleted of chemokines critical for DC and T cell recruitment.

Here we demonstrated that *Keap1*-mutant tumors suppress CD8, but not CD4, T cell immunosurveillance promoting CD8 T cell exhaustion characterized by reduced expression of activation markers, decreased proliferative capacity and production of effector molecules. We also showed that T lymphocytes and dendritic cells infiltrating KP tumors express receptors for chemokines robustly downregulated in *Keap1*-mutant tumor cells. Taken together, our characterization of the immune landscape in *Keap1* wild-type and mutant tumors and the outlined depletion studies provide critical insight into how *Keap1* mutation interferes with the cDC1 – CD8 anti-tumor axis, thereby disrupting immune surveillance.

### Keap1 mutation impairs response to immune checkpoint blockade

Engagement of cDC1 and T cells is a critical determinant of success of immunotherapies in several preclinical models of cancer (18-21). Moreover, recent evidence implicates tumor intrinsic mutations in *KEAP1* with immunotherapy resistance in humans (33,34). This prompted us to investigate whether suppression of DC-CD8 T cell axis by *Keap1*-mutant tumors can drive resistance to checkpoint blockade.

Despite several efforts to establish genetically-defined KRAS-driven LUAD mouse models that induce T cell responses and can respond to checkpoint blockade therapy (38,39,42,66-68), to the best of our knowledge, none have been shown to respond to checkpoint blockade therapy (40,42). Here we used our H-Y-driven orthotopic mouse model to study how *Keap1*-mutation subverts anti-tumor immune responses to checkpoint inhibition.

We wanted to investigate immune responses and tumor growth kinetics of wild-type and mutant tumors following immunotherapy. We injected wild-type and mutant male KP cell lines orthotopically into C57BL/6J female mice (Fig. 4A). After validating tumor engraftment using *in vivo* bioluminescence imaging, we treated mice with anti-PD1 monoclonal antibody (clone 29F.1A12). We selected this clone following rigorous testing of different anti-PD1 clones in an MC38 colon adenocarcinoma model which revealed dramatic differences in efficacies of the 3 widely used anti-PD1 clones in respect to tumor growth (Fig. S5A-B). It has been shown that higher tumor burden can promote resistance to checkpoint blockade (69). For this reason, we initiated anti-PD1 treatment when tumor burden was equal between the two genotypes (Fig S5C). Wild-type tumors responded while anti-PD1 treatment had no effect on the growth of *Keap1*-mutant tumors (Fig. 4A-B). Specifically, half (3/6) of the mice with *Keap1* WT tumors showed robust regression, 2/6 showed initial regression followed by growth, and tumor growth in 1/6 was stabilized. On the other hand, none of the *Keap1*-mutant tumors displayed sustained regression; 2/8 mice initially regressed but then started growing soon after, growth in 2/8 mice stabilized and growth in 4/8 mice continued unperturbed (Fig. S5D). These results are consistent with observations from clinical studies that suggest a central role of KEAP1 in resistance to checkpoint inhibition(33,34) and confirm that *Keap1* mutation in the tumor cells promotes an immunosuppressive microenvironment driving resistance to checkpoint blockade therapy.

**Figure 4:**
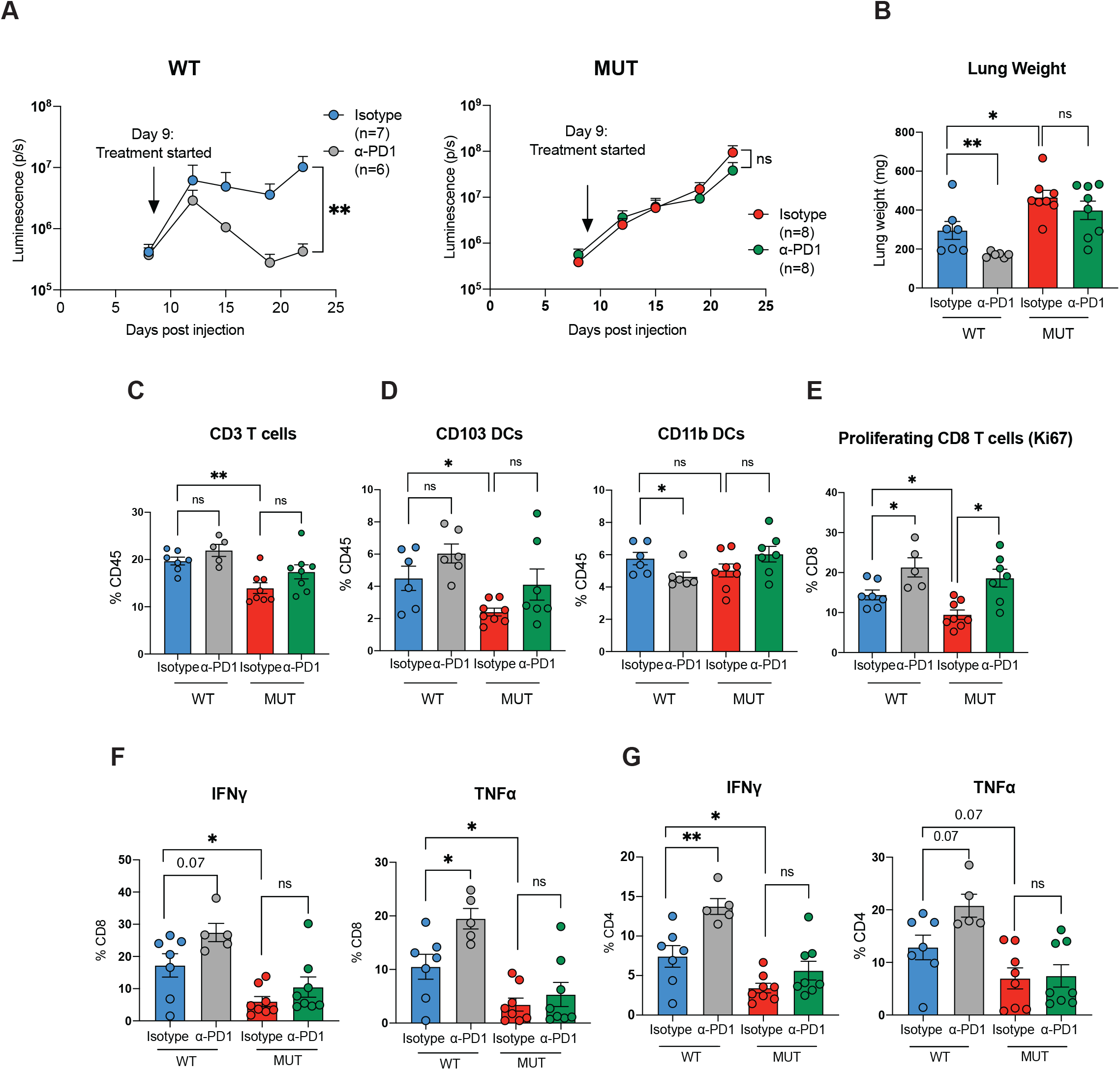
*Keap1* mutation drives immunotherapy resistance in antigenic Kras-driven lung adenocarcinoma mouse model. (A) Growth kinetics of *Keap1* wild-type (left) and mutant (right) KP cells injected i.v. in female C57BL/6J hosts upon treatment with anti-PD1 monoclonal antibody or isotype control. Treatment was initiated 9 days post-injection and continued until experimental endpoint. Each experimental subgroup had n ≥ 6 mice. Tumor engraftment was confirmed in all mice prior to randomization into anti-PD1 and isotype treatment groups. (B) Lung weight as proxy for tumor burden measured on day 24 in mice bearing *Keap1* wild-type and mutant tumors. Each experimental subgroup had n ≥ 6 mice. (C) Percentage of CD3+ lymphocytes among tissue-infiltrating immune cells. Each symbol represents an individual mouse. Each experimental subgroup had n ≥ 5 mice. (D) Percentage of CD103+ and CD11b+ DCs among tissue-infiltrating immune cells in the lungs of animals with *Keap1* wild-type and mutant tumors treated with anti-PD1 or isotype control. Each symbol represents an individual mouse (E) Percentage of proliferating (Ki67+) cells within the CD8+ T cell gate in *Keap1* wild-type and mutant tumor bearing mice treated with anti-PD1 or isotype control. Each symbol represents an individual mouse. Each experimental subgroup had at least n=5 mice. (F) Percentage of intracellular IFNγ+ and TNFα+ cells among the CD8+ T lymphocytes within wild-type and mutant *Keap1* tumors treated with anti-PD1 or isotype control. Each symbol represents an individual mouse. Each experimental subgroup had n ≥ 5 mice (G) Percentage of intracellular IFNγ+ and TNFα+ among the CD4+ T cells in the wild-type and mutant *Keap1* tumors treated with anti-PD1 or isotype control. Each symbol represents an individual mouse. Each experimental subgroup had n ≥ 5 mice. *P<0.05; **P<0.01; ***P<0.001; ****P < 0.0001

We then sought to determine key immunotherapy response readouts by evaluating changes in the immune landscape. We did not observe changes in the numbers of T lymphocytes or DC populations (Fig. 4C-D) upon anti-PD1 treatment. However, immunotherapy treatment resulted in increased CD8 T cell proliferation irrespective of *Keap1* status (Fig. 4E). We then investigated the production of effector molecules by T lymphocytes. We observed increased expression of IFN-γ and TNF-α in both CD4 and CD8 T cells in *Keap1* wild-type but not mutant tumors (Fig. 4F-G). These results indicate that although antibody treatment promotes proliferation of CD8 T cells in both genotypes, anti-tumor T cell responses are blunted in *Keap1-*mutant tumors, while in WT tumors T cell response is characterized by upregulation of effector molecules essential for tumor control.

### Keap1 mutation correlates with exclusion of DCs and CD8 T cells in human lung adenocarcinoma

We next examined whether human tumors with KEAP1 mutation display signs of immune evasion that can drive resistance to checkpoint blockade. To evaluate this, we assessed the effect of *KEAP1* mutation on the immune microenvironment of primary human LUAD tumor samples. We previously identified *KEAP1* mutant LUAD tumors using targeted exome capture and validated these results via NQO1 staining (31). Using multiplex immunofluorescence, we found that *KEAP1* wild-type tumors have increased infiltration of total CD3 T cells, CD8 T cells and PD1-expressing CD8 T cells compared to mutant tumors (Fig. 5A-B; Fig. S6A). Our mouse data suggest that *Keap1*-mutant tumors do not appear to suppress CD4 T cells responses (Fig. S4B-C; S4G-H). In line with results from our mouse models of this malignancy, we did not observe any significant differences in the CD4 T cell infiltration patient samples irrespective of their *KEAP1* mutation status (Fig. S6B).

**Figure 5:**
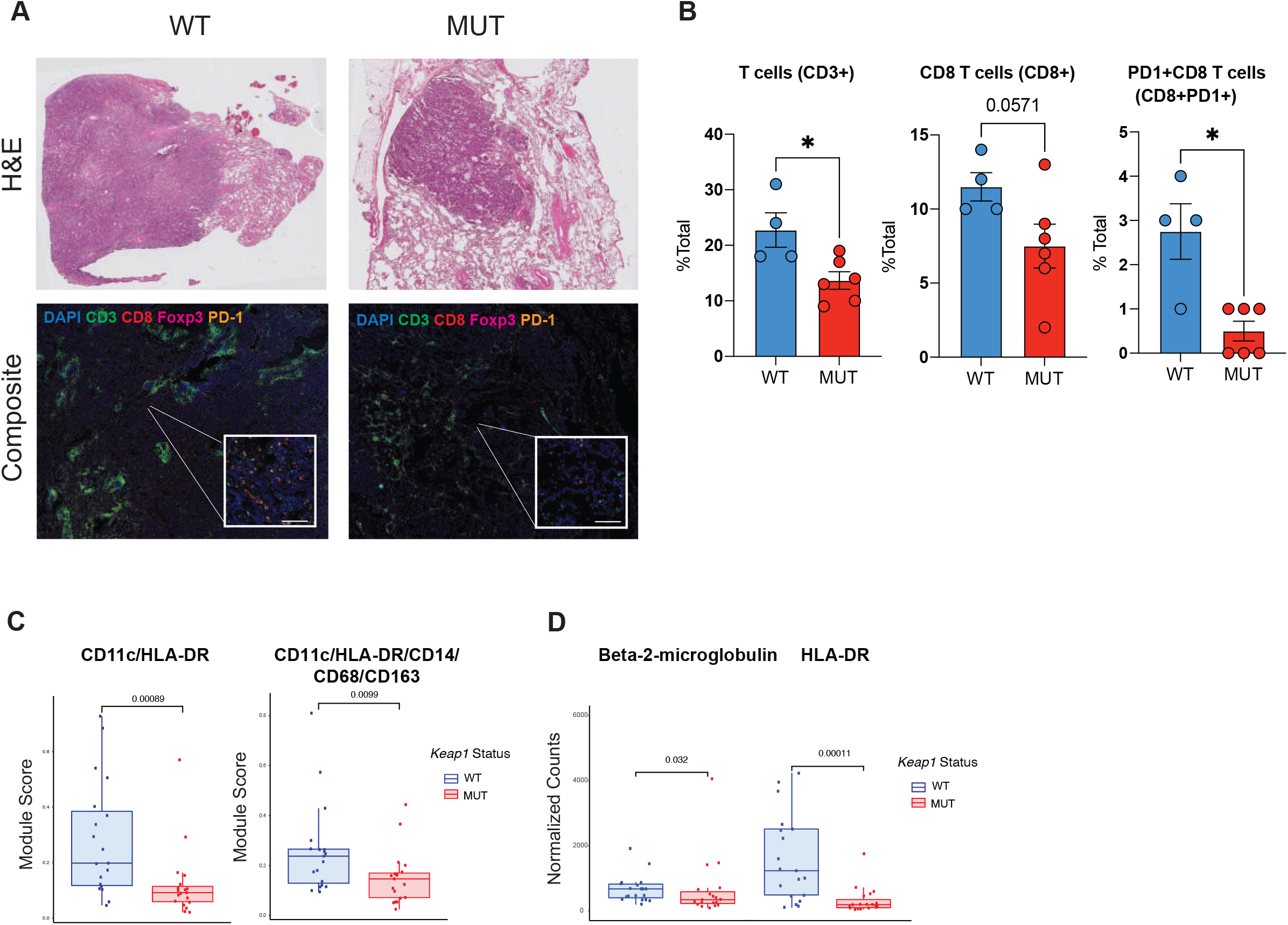
Genetic inactivation of *KEAP1* drives immune evasion in human LUAD tumors. (A) Representative H&E (top) and immunofluorescence (bottom) images of *KEAP1* wild-type and mutant human tumors. DAPI is shown in blue, CD3 in green, CD8 in red, Foxp3 in magenta, PD1 in orange. Scale bar 20um. (B) Quantification of T cells (CD3+), CD8 T cells (CD8+) and PD1+ CD8 T cells (CD8+PD1+) in *KEAP1* wild-type and mutant human tumors. Tumor area was identified based on H&E staining. (C) Box plots showing expression of CD11c/HLA-DR and CD11C/HLA-DR/CD14/CD68/CD163 protein modules in *KEAP1* wild-type (n=19) and mutant (n=19) patient tumor samples from tumor microarray containing LUAD patient samples. Microarray was stained and analyzed using Nanostring GeoMx platform. Expression is shown for pancytokeratin positive areas. (D) Box plots showing expression for individual proteins: beta-2-microglobulin, HLA-DR and Sting. *P<0.05; **P<0.01; ***P<0.001; ****P < 0.0001

To assess infiltration of antigen presenting cells (DCs) we used Nanostring GeoMx digital spatial profiling to perform high-plex proteomic analysis with spatial resolution. Using a tissue microarray (TMA) containing Keap1 wild-type and mutant LUAD human samples, we acquired GeoMx data from three different areas of each sample (Fig. S6C). We developed protein modules that describe DCs (CD11c/HLA-DR) and mononuclear phagocytes (CD11c/HLA-DR/CD14/CD68/CD163) and evaluated the TMAs in which we distinguished between tumor and adjacent normal tissue based on pankeratin staining (Fig. 5C, Fig. S6C). We observed a reduction in the expression of both modules in *KEAP1*-mutant tumors compared to patient samples in which the allele had no evident mutations (Fig. 5C), consistent with our pre-clinical mouse model data. Interestingly, *KEAP1* mutation status negatively correlated with levels of proteins involved in antigen presentation (beta-2-microglobulin, HLA-DR) (15,70) (Fig. 5D). This suggests that genetic inactivation of KEAP1 is a mechanism co-opted by tumor cells to avoid immunosurveillance in both mice and humans.

## Discussion

Cancer immune evasion is a major obstacle in designing effective anticancer therapies (71). Immunoediting, a process by which less immunogenic tumor cells are selected over time, contributes to evasion of immune surveillance (71-74). Reduced immunogenicity can be a consequence of diminished antigen presentation, decreased recruitment and activation of T cells, or a result of increased recruitment of suppressive cells or production of immunosuppressive factors.

There is accumulating evidence that tumor cell-intrinsic alterations in signaling pathways, previously described to drive tumorigenesis, can promote cancer immune evasion via modulation of TIME (75). Tumor-intrinsic alterations can promote a “cold” tumor microenvironment via exclusion of cells essential for a productive immune response or via attraction of immunosuppressive populations (75). Cancer genome sequencing studies have revealed that *KEAP1* is frequently mutated in NSCLC (23). Emerging clinical data highlights that alterations in *KEAP1* are associated with poor prognosis and resistance to multiple therapies including immunotherapy (33,34,76). Interestingly, *KEAP1*-mutant tumors have also been found to be associated with a “cold”, diminished of T cells phenotype (35). Understanding the underlying mechanisms of KEAP1-dependent immune evasion can pave the way to new immune-based therapeutic interventions. Here we demonstrated that *Keap1* mutation suppresses cDC1 orchestrated CD8 T cell immunity and drives immunotherapy resistance. Our findings suggest that genetic inactivation of KEAP1 in LUAD tumors could function as fundamental immune evasion mechanism.

We demonstrate that there is no difference in growth kinetics of *Keap1*-mutant and wild-type tumors in immunodeficient hosts or in tumor models lacking antigenicity (Fig. 1G-I). This indicates that a major mechanism by which *Keap1* mutation drives accelerated tumor growth *in vivo* is via promoting immune evasion. Our findings from the orthotopic tumor models rely on transplantation of cell lines and thus tumor engraftment could be contributing to the observed differential growth *in vivo*. However, we initiated our depletion studies only once engraftment was validated and randomized our cohorts to ensure comparable tumor burden at the beginning of our studies. Furthermore, our results from autochthonous models recapitulated these findings and analysis of human samples demonstrate that tumor-intrinsic *KEAP1* mutation promotes immune evasion in cancer patients as well.

T cell infiltration in the tumor microenvironment has been correlated with better prognosis and responses to checkpoint blockade (9,77). Effective T cell infiltration involves both cytotoxic CD8 T cells and IFNγ producing Th1 CD4 T cells (12,14,78,79). Although dendritic cells were primarily studied for their ability to prime T cells in the lymph nodes, they are now also recognized as key players at primary tumor site (13,19,61). DCs can take up dead tumor cells and present internalized antigens on major histocompatibility complex (MHC) molecules. Peptides loaded on MHC-I molecules are recognized by antigen-specific CD8 T cells leading to their activation, proliferation and up-regulation of cytotoxic molecules. Here we demonstrated that *Keap1*-mutant lung tumors are characterized by decreased cDC1 infiltration and activation (Fig. 1E-H; Fig. S3B), resulting in diminished CD8 effector T cell responses (Fig. 3H-I). Our analysis reveals that CD8 T cells from *Keap1*-mutant tumors appear more exhausted, have lower levels of activation markers, decreased proliferative capacity and reduced production of effector molecules (Fig. 3A-G).

We demonstrate that *Keap1* genetic inactivation is associated with decreased type I/II interferon signatures and chemokine expression (Fig.1J-K), previously shown to be essential for activation of DC-T cell immune axis (12-14). Additional studies are needed to fully elucidate the role of individual chemokines and interferons in KEAP1-mediated immunosuppression as well the dynamic interaction of DCs and T cells in wild-type (DC-high) compared to mutant (DC-low) TIME. Finally, *Keap1*-mutant tumors could potentially affect other immune populations. For instance, we observed a decrease in Th1 cells with contemporaneous increase in T regulatory cells in mutant tumors (Fig. S3D,F). We speculate that *Keap1* mutations could be differentially impacting both anti-tumor and tumor-promoting immune populations.

Since *Keap1*-mutant tumors drastically modulate the TIME via suppressing cDC1 and CD8 T cell responses essential for immunotherapy responses, we wanted to investigate whether *Keap1* mutation could be driving resistance to checkpoint blockade. There have been multiple efforts to develop genetically defined syngeneic KRAS-driven LUAD mouse models that are able to induce T cell responses and respond to checkpoint blockade therapy. Most of immunogenic cancer models rely on the expression of model antigens, such as ovalbumin and LCMV derived peptides (38,39,66,67), or mismatch repair genetic variants (42,68). Importantly, none of these models have been shown to respond to checkpoint blockade therapy (40,42) hindering studies aimed at investigating immunotherapy responses. We hypothesized that the lack of responses might be driven by two main factors; the absence of a diverse antigenic landscape and/or inefficient treatment regimens. After validating that our H-Y-driven antigenic model can drive anti-tumor T cell responses (Fig. 1B-D), we sought to optimize checkpoint blockade treatment protocols. Recent work suggested that some anti-PD1 clones widely used in preclinical models, including LUAD models, deplete antigen specific T cells (80). For this reason, we tested multiple anti-PD1 clones on a subcutaneous colon cancer model that robustly responds to checkpoint therapy to select an anti-PD1 clone that reliably drives tumor regression (Fig. S5). We used our antigenic H-Y-driven model with appropriate anti-PD1 treatment regime and demonstrated that *Keap1* wild-type tumors respond to immunotherapy while *Keap1*-mutant tumors fail to respond to checkpoint-inhibitor treatment.

We showed that tumor-intrinsic *Keap1* mutation results in downregulation of interferon pathways and chemokine expression. NRF2, the most well characterized target of KEAP1, has been suggested to transcriptionally regulate immunomodulatory pathways such as chemokine production and cGAS/STING-induced type I interferon pathway, albeit primarily in an *in vitro* or non-tumor setting (81-83). Given that point mutations within *KEAP1* impair targeting and degradation of multiple ETGE motif containing substrates (PALB2, MCM3, IKKB, DPP3) (84-87), not only of NRF2, it remains to be determined whether it is NRF2 or other substrates underlie the immune evasion phenotype of KEAP1-mutant tumors.

Finally, our work has important clinical implications. We revealed that a major genetic subset of LUAD patients, patients with *KEAP1* mutation, is characterized by an immunosuppressive phenotype (Fig. 2-3,5) and is resistant to checkpoint blockade (Fig. 4). Our group has previously showed that *Keap1*-mutant tumors are highly sensitive to glutaminase inhibitor CB-839 (31,88,89) which blocks the conversion of glutamine to glutamate. Modulating glutamine metabolism was previously shown to, not only inhibit tumor growth, but also promote restoration of anti-tumor immunity (90). Importantly, glutaminase inhibition with CB-839 is currently being tested in phase 2 clinical trials in *KEAP1* or *NRF2-*mutant LUAD patients as a single agent (NCT03872427) or in combination with standard of care checkpoint inhibition and chemotherapy (KEAPSAKE: NCT04265534). We hope our studies will encourage development of new therapeutic strategies combining glutamine metabolism inhibition with checkpoint blockade therapies to improve immunotherapy responses in *KEAP1*-mutant tumors.

## Supporting information

All supplemental and legends

## Author Contributions

A.M.Z. conducted experiments with assistance from R.P, T.K, A.R., S.S., M.C. A.M.Z, Y.H. and T.B. did the bioinformatic analysis. W.L.W generated the Keap1 vectors. A.M.Z and E.B carried out mouse immunofluorescence. A.M.Z. and A.H. conducted the scRNA-seq. A.K. provided advice and reagents for checkpoint blockade experiments. R.P and A.Y helped with Nanostring analysis. E.I. advised on initial flow cytometry experiments. C.W, J.P, C.R provided human tissue microarray. A.L.M. provided human samples and advice. K.M.K. supervised the mouse immunofluorescence experiments. A.T. supervised the bioinformatics analyses. K.K.W. provided conceptual advice. A.M.Z., T.P., S.BK. conceived the study, designed the experiments and wrote the manuscript. All authors reviewed and discussed the final version of the paper.

## Acknowledgements

Work in T.P. laboratory was supported by NIH grants (R37CA222504, R01CA227649) and an American Cancer Society Research Scholar Grant (RSG-17-200-01-TBE). Work in S.B.K. laboratory was supported by NIH grants (R01HL-125816 and R03AI144714), LEO Foundation Grant (LF-OC-20-000351), NYU Cancer Center Pilot grant (P30CA016087), and the Judith and Stewart Colton Center for Autoimmunity Pilot grant. W.L.W is supported by NIH training grants (T32GM007308 and 1F30CA247020). R.G.P is supported by NIH training grant (T32CA009161). E.B. is supported by NIH training grant (T32AI100853-10). Work in K.M.K. laboratory is supported by NIH grants (R01AI143861-01). A.T. is supported by the NCI/NIH P01CA229086 and NCI/NIH R01CA252239. T.B. was supported by the Danish Cancer Society (Kræftens Bekæmpelse). We thank the NYU Langone Experimental Pathology Laboratory, Flow Cytometry Core and Animal Resources Facility staff for their support and guidance. We would like to thank the Genome Technology Center (GTC) for expert library preparation and sequencing, and the Applied Bioinformatics Laboratories (ABL) for providing bioinformatics support and helping with the analysis and interpretation of the data. GTC and ABL are shared resources partially supported by the NYU Cancer Center Support Grant P30CA016087. This work has used computing resources at the NYU School of Medicine High Performance Computing Facility.

## MATERIALS AND METHODS

### Mice

All animal procedures were approved by the NYU Langone Medical Center Institutional Animal Care and Use Committee (IACUC). Animals were housed according to IACUC guidelines in ventilated caging in a specific pathogen free (SPF) animal facility. C57BL/6J mice were bred in house or purchased from Jackson Laboratories. B6.129S(C)-Batf3^tm1Kmm^/J were purchased from Jackson Laboratories. Animals acquired from Jackson laboratories were housed for at least one week in our SPF facility prior to experiment initiation. *Kras*^LSL-G12D/+^; *p53*^fl/fl^ were crossed to *Keap1*^fl/fl^ (Taconic, 8799). For tumor induction, mice 6-10 weeks of age were infected intratracheally with Cre-expressing lentivirus as previously described (91). Males or female mice were used as specified in the figure legends.

### Tumor Cell Line Generation

*Kras*^LSL-G12D/+^; *p53*^fl/fl^ cells were isolated from a male mouse. No clonal selection was performed. We generated *Keap1* overexpressing cells by cloning the mouse *Keap1* cDNA into Gibson compatible lentiviral backbone with hygromycin resistance cassette. *Keap1*R470C point mutation was generated by QuickChange II Site-directed Mutagenesis (Agilent) per manufacturer’s protocol. Cells infected were selected with 600μg/ml hygromycin. All cell lines tested negative for mycoplasma (PlasmoTest, InvivoGen). Cells were maintained in DMEM (Cellgro, Corning) supplemented with 10% fetal bovine serum (FBS) (Sigma Aldrich) and gentamycin (Invitrogen).

### Tumor Cell Injections

Tumor cells were harvested by trypsinization and washed three times with PBS. For lung orthotopic experiments, 7-10-week-old mice were injected intravenously with 2×10^5^ KP cells expressing GFP and luciferase in 200ul endotoxin-free PBS. For MC38 subcutaneous injections, cells were injected into both right and left flack of each recipient mouse at 5×10^5^ cells in 100ul of endotoxin free PBS containing 2.5mM EDTA.

### Virus preparation

Lentivirus was produced by co-transfection of HEK293 cells with viral vector and packaging plasmids (psPAX2, pMD2G) using PEI Pro (PolyPlus). Virus containing media was collected 48h and 72h post transfection and filtered through 0.45uM PVDF filter. For cell line generation, harvested virus was added onto target cells in the presence of 8μg/ml polybrene (Millipore). For *in vivo* experiments, harvested virus was concentrated by ultracentrifugation at 25000rpm for 2h ate 4°C, dissolved in PBS and stored at −80°C. Virus titration was performed using the Green-Go reporter cell line as previously described (92).

### Proliferation Assay

For cell proliferation assays, cells cultured in DMEM/10% FBS were trypsinized, and plated into 6 well plates at 2*10^5 cells and 2ml DMEM/10%FBS media per well. Cells were counted at days 1, 3 and 6 post plating.

### UTY qPCR

mRNA was harvested from tumor cells using RNeasy mini kit (Qiagen) according to manufacturer’s protocol. cDNA was synthesized from mRNA using SuperScript VILO (Invitrogen) according to manufacturer’s protocol. Real-time PCR was performed with StepOne Plus PCR system using SybrGreen master mix (Applied Biosystems). Primer sequences used were as follows: UTY forward, 5’-GAGGTTTTGTGGCATGGGAG -3’; UTY reverse, 5’-TGCAGAAGATAACGAAGGAGCTA-3’

### Antibody depletion experiments

For depletion of CD8+ and CD4+ T lymphocytes, anti-CD8a antibodies (clone: 2.43, Bioxcell) and anti-CD4 (GK1.5, Bioxcell) were used. Antibodies were diluted in PBS and injected intraperitoneally at 150μg/mouse twice a week (Monday, Friday), 7 days after tumor cell implantation until the end point of the experiment.

### Labeling of Circulating Cells

Circulating immune cells were labeled using intravenously injected APC-conjugated CD45 antibody (30-F11, Biolegend) 3 minutes prior to sacrifice as previously described (39,93).

### Histology analysis

Mice were euthanized by lethal doses of ketamine and xylazine. Lungs were inflated and incubated overnight at room temperature (RT) with 10% formalin. They were subsequently transferred to 70% ethanol, and subsequently embedded in paraffin. 5uM sections were stained with H&E or subjected to other immunohistochemical staining. For immunohistochemistry (IHC) we used antibody against Nqo1 (1:100, HPA007308, Sigma-Aldrich). Immunohistochemitry was performed on a Leica Bond RX and slides were imaged on a Leica SCN400F whole slide scanner. For Nqo1, antigen retrieval was performed using antigen retrieval buffer pH=6 (Leica) for 20min. For detection, Leica Bond Polymer Refine Detection secondary antibody was used according to manufacturer’s protocol. Total tumor lung area was quantified via H&E-stained slides using QuPath software (94). Tumor burden and IHC analyses were done in blinded fashion.

### Immune Cell isolation from lung tumors

Lungs were harvested at timepoints indicated in figure legends. Lungs were minced on glass slides followed by digestion using a cocktail of collagenase IV (125U/ml, Sigma), DNASE I (40U/ml, LifeTechnologies), 1X HEPES (Cellgro) in RPMI. Samples were incubated at 37°C for 35min and mixed by inverting every 5-8min. Digestion was quenched by adding RPMI containing 1mM EDTA final concentration. Digested samples were strained through 70μm cell strainers followed by red blood cell lysis (BD PharmLyse, BD Biosciences) for 10min at 4°C.

### Checkpoint blockade treatments

For checkpoint blockade treatment optimization on mc38 tumors anti-PD1 (RMP1-14, Bioxcell), anti-PD1 (mPD1-4H2-mIgG1-D265A, clone 6A1_RAS_Ab, Bristol Myers Squibb), anti-PD1 (29F.1A12, Bioxcell), anti-PD-L1 (10F.9G2, Bioxcell) was used. For isotype controls, IgG2b (LTF-2, Bioxcell), IgG1 (Bristol Myers Squibb), IgG2a (2A3, Bioxcell) were used. Tumor volume was measured by caliper, with volume calculated based on the following formula: (Length x Width^2 x (3.14/6)). After tumor engraftment (∼40-50mm^3^ tumor size) and before tumors reach 100mm^3^, animals were randomized and assigned to treatment group. For treatments of KP lung orthotopic tumors, anti-PD1 (29F.1A12, Bioxcell) was used. For anti-PD1 treatments, antibodies were diluted in PBS and injected intraperitoneally at 200ug/mouse three times a week until the end point of the experiment. For anti-PD-L1 treatment, mice were given a total of three doses intraperitoneally at 200μg/mouse every other day.

### Mouse Tissue Immunofluorescence

Tissues were processed as previously described (95). Harvested tissues were fixed in PLP buffer (96) overnight at 4°C. Tissues were then dehydrated in 30% sucrose for 24 hours at 4°C and subsequently embedded in OCT media and stored at −80°C. 20μm sections were loaded on to racks (Shandon Sequenza) and washed with PBS. Sections were blocked with anti-CD16/CD32 (BioLegend, clone 93, 1/300) diluted in 1x PBS, 2% FBS, and 2% goat serum for 1h at RT. Staining antibodies were diluted in 1x PBS with 2% FBS, and 2% goat serum. Following blocking, sections were then incubated with fluorescently conjugated antibodies for CD11c (N418, Biolegend, 1/100), CD4, (Gk1.5, Biolegend,1/100), CD103 (M290, Biolegend, 1/100) diluted in PBS with 2%FBS and 2% goat serum for 1h at room temperature. Zeiss LSM 880 confocal microscope with the Zeiss Zen Black software was used for imaging. Confocal microscopy images were analyzed using the Bitplane Imaris x64 version 9.0.2 software. Images were filtered with the 3×3×1 median filter function to reduce background autofluorescence. To quantify different cell types within tumor lesions, “spots” were made from either the positive or negative signal combinations of CD45circ-CD11c+ CD103+ (cDC1). Tumors were manually constructed into “2D surfaces” based on positive GFP signal. The “find spots close to surfaces” function was then performed on generated spots and surfaces with a threshold of 0.5 to quantify generated spots located within the surfaces. These quantifications are reported as the number of spots present within constructed tumor surfaces per surface area to control for differences in tumor area between images.

### Human Tissue Immunofluorescence

5μm paraffin sections were stained using Akoya Biosciences® Opal™ multiplex automation kit reagents unless stated otherwise. Automated staining was performed on Leica BondRX® autostainer. The protocol was performed according to manufacturers’ instructions with the following antibodies: anti-CD3 (LN10, Biocare Medical, 1/75, Opal570), anti-CD8 (C8/144, Cell Signaling Technology, 1/500, Opal620), anti-PD-1 (D4W2J, Cell Signaling Technology, 1/200, Opal520), anti-Foxp3 (236A/E7, Thermo Fisher Scientific, 1/100, Opal780). All slides underwent sequential epitope retrieval with Leica Biosystems epitope retrieval 2 solution (ER2, EDTA based, pH9, Cat. AR9640), primary and secondary antibody incubation and tyramide signal amplification (TSA) with Opal® fluorophores. Primary and secondary antibodies were removed during epitope retrieval steps while fluorophores remain covalently attached to the epitope. Semi-automated image acquisition was performed on a Vectra® Polaris multispectral imaging system. After whole slide scanning at 20X the tissue was manually outlined to select fields for spectral unmixing and analysis using InForm® version 2.4.7 software from Akoya Biosciences. For each field of view, a tissue segmentation algorithm (tissue/no tissue) was run prior to cell segmentation. Cells were segmented based on nuclear signal (DAPI). Cells were phenotyped after segmentation using inForm’s trainable algorithm based on glmnet1 package in R. Phenotypes were reviewed for different samples during training iterations.

### Flow Cytometry and FACS

For surface staining, single cell suspensions were incubated for 10min with Fc receptor block (2.4G2, Bioxcell) at 4°C, followed by antibody staining for either 15min (for adaptive immune markers) or 30min (for innate immune markers) at 4°C. Staining was performed in FACS buffer (PBS, 2%BSA, 1mM EDTA). For cytokine staining, single cell suspensions were treated with a cell stimulation and protein transport inhibition cocktail, containing Golgi Plug (55029, 1/1000, BD Biosciences), Golgi Stop (555029, 1/1000, BD Biosciences), PMA (1/10000), Ionomycin (1/1000) for 3.5h. at 37°C in RPMI supplemented with 10%FBS. Cells were then surface stained as above, fixed in 2%PFA and permeabilized with 0.5% saponin. For transcription factors, staining was performed with the Foxp3 Staining Buffer kit (00552300, eBioscience). Fluorochrome conjugated antibodies with the following specificities were used: CD3 (ebio500A2, ebioscience), CD4 (RM4-5, ebioscience), CD4 (GK1.5, BD bioscience), CD45 (30-F11, BD bioscience), CD8 (53-6.7, BD bioscience), CD44 (IM7, ebioscience), CD62L (MEL-14, ebioscience), CD69 (H1.2F3, ebioscience), Gr-1 (RB6-8C5, ebioscience), Rorgt (Q31-378, BD Bioscience), Ki67 (B56, BD Bioscience), IFN-γ (XMG1.2, ebioscience), TNF-α (MP6-XT22, ebioscience), Foxp3 (FJK-16s, ebioscience), Tbet (4B10, ebioscience), GATA3 (TWAJ, ebioscience), TCRb (H57-597, ebioscience), PD-1 (J43, ebioscience), CD86 (GL1, BD Bioscience), CD11b (M1-70, Biolegend), CD11c (N418, Biolegend), CD103 (2E7, Biolegend), CD64 (X54-5/7.1, Biolegend), CD80 (16-10A1, Biolegend), IA/IE (M5/114.15.2, eBioscience), Siglec-F (E50-2440, BD Bioscience). For viability dye (1/1000, Biolegend), cells were stained in 100ul PBS for 10min at RT. Samples were acquired using BD LSR Fortessa Cell Analyzer For sorting, single cell suspensions were resuspended in FACS buffer (PBS, 2%FBS, 0.01%Tween) and stained for viability dye (1/1000, Biolegend) in 100ul PBS for 10min at RT followed by surface staining with antibody specific for CD45 (30-F11, PercPcy55, 1/300, Biolegend). Tumor cells (for RNA-seq) were sorted for singlets, live cells, negative for CD45 and GFP positive. Immune cells (for single cell RNA seq) were sorted for singlets, live cells, negative for CD45 circulating antibody, and CD45 positive. Cells were sorted into pre-chilled tube containing DMEM supplemented with 20% FBS. Data were analyzed with FlowJo software (Tree Star Inc).

### RNA-seq

Tumor cells were sorted as described above. RNA was isolated using Purelink RNA mini kit (Invitrogen) per manufacturer’s protocol. cDNA was synthesized from RNA using SMARTer PCR cDNA synthesis kit (Clontech) per manufacturer’s protocol. Sequencing libraries were prepared using Nextera XT DNA library preparation kit (Illumina) per manufacturer’s protocol. Samples were pooled at equimolar ratios. Libraries were loaded on an SP11 cycle flow cells and sequenced on Illumina NovaSeq6000. Read qualities were evaluated using FASTQC (Babraham Institute) and mapping to GRCm38 (GENCODE M25) reference genome using STAR programs (97) and RSEM (98) and with default parameters. Read counts, TPM and FPKM were calculated using RSEM. Identification of differentially expressed genes between wild-type and mutant tumor cells was performed using DESeq2 in R/Bioconductor. All plots were generated using customized R scripts. Pathway enrichment analysis was performed using GSEA program (99) based on log2FC values. Gene sets were downloaded from MsigDB (100).

### Single cell RNA seq

Samples were multiplexed using cell hashing antibodies. These were covalently and irreversibly conjugated to barcode oligos by iEDDA-click chemistry (“home conjugated”) as previously described (101,102). Cells were incubated for 10min with Fc receptor block (2.4G2, Bioxcell) and subsequently with hashing antibodies for 30min at 4°C. Cells were washed three times in PBS containing 2% bovine serum albumin (BSA) and 0.01% Tween followed by centrifugation (300g) for 5min at 4°C and supernatant aspiration. After final wash, cells were resuspended in PBS and filtered through 40μm cell strained. Cells from each sample were pooled and loaded into 10X Chromium. Gene expression together with Hashtag oligo (HTO) libraries were processed using Cell Ranger (v5.0.0) in count mode. UMI count matrices from each modality were imported into the same Seurat object as separate assays. Cell-containing droplets were selected using the default filtering from Cell Ranger count “filtered_feature_bc_matrix”. Viable cells were filtered based on having more than 250 genes detected and less than 55% of total UMIs stemming from mitochondrial transcripts. HTO counts were normalized using centered log ratio transformation before hashed samples were demultiplexing using the Seurat::HTODemux function (positive.quantile set to 0.93). Cells from two separate droplet emulsion reactions were combined using the standard SCTransform integration workflow in the Seurat (v4) R package (103). Multimodal integration was performed using the weighted-nearest neighbor (WNN) method available in Seurat. Briefly, a WNN network was constructed based on modality weights estimated for each cell using FindMultiModalNeighbors function. Cell clusters were identified using FindClusters function based on the weighted SNN (WSNN) graph using leiden algorithm at various resolutions, ranging from 1 to 1.6. Cell types were annotated based on canonical cell type markers as well as differential expressed genes of each cluster identified using FindAllMarkers function in Seurat. Clusters expressing markers of the same cell type were merged into a single cluster.

### Nanostring

Slide preparation was performed following the GeoMx DSP instructions. Briefly, 5μm TMA sections were baked for 2 hrs at 60 degrees Celsius before loading onto an automated slide stainer (LeicaBiosystems Bond RX). Slides underwent Baking and Dewaxing steps followed by blocking for 60 minutes with buffer ‘W’ (part of the slide prep kit for GeoMx). GeoMx antibody incubation was performed overnight at four degrees Celsius with the following combination of antibody panels: Human Immune Cell Profiling Protein Core, Human Immune Activation Status Protein, Human Immune Cell Typing Protein, Human IO Drug Target Protein, Human Pan-Tumor Protein (v1.0 for all panels and core). The Human Solid Tumor TME (v1.0) morphology marker kit was used to visualize pancytokeratin and CD45 antigens on the slide scan. Digital counts from barcodes corresponding to protein probes were analyzed as follows: raw counts were first normalized with internal spike-in controls (ERCC) to account for system variation. Data was normalized using the geometric mean of housekeeping antibody counts for Histone H3 and S6. The module score was calculated by first rescaling each gene across measurements by dividing by the maximum value across measurements, then for each measurement, taking the mean of genes included in each module. Normalized values were visualized with boxplots. The wilcoxon test was used to calculate statistics between conditions. One outlier sample was removed. Outlier samples were determined as having values above the 98^th^ percentile in more than 20 proteins. Ggplot and ggpubr R packages were used to generate plots and statistics.

### Statistics

Values are represented as mean ± SEM. Statistical analyses were performed using Prism 9 (GraphPad Software) and a p value<0.05 was considered significant. All animal experiments contained at least n>3 mice. Mann-Whitney test for flow cytometry and lung weight experimemts with less than 2 experimental groups. One-way analysis of variance (ANOVA) and Tukey’s test for lung weights and histological analysis with more than two experimental groups. Two-way ANOVA for growth *in vivo* growth kinetics experiments. *P<0.05; **P<0.01; ***P<0.001; ****P < 0.0001

