## Supplementary material for "KEAP1 mutation in lung adenocarcinoma promotes immune evasion and immunotherapy resistance": All supplemental and legends

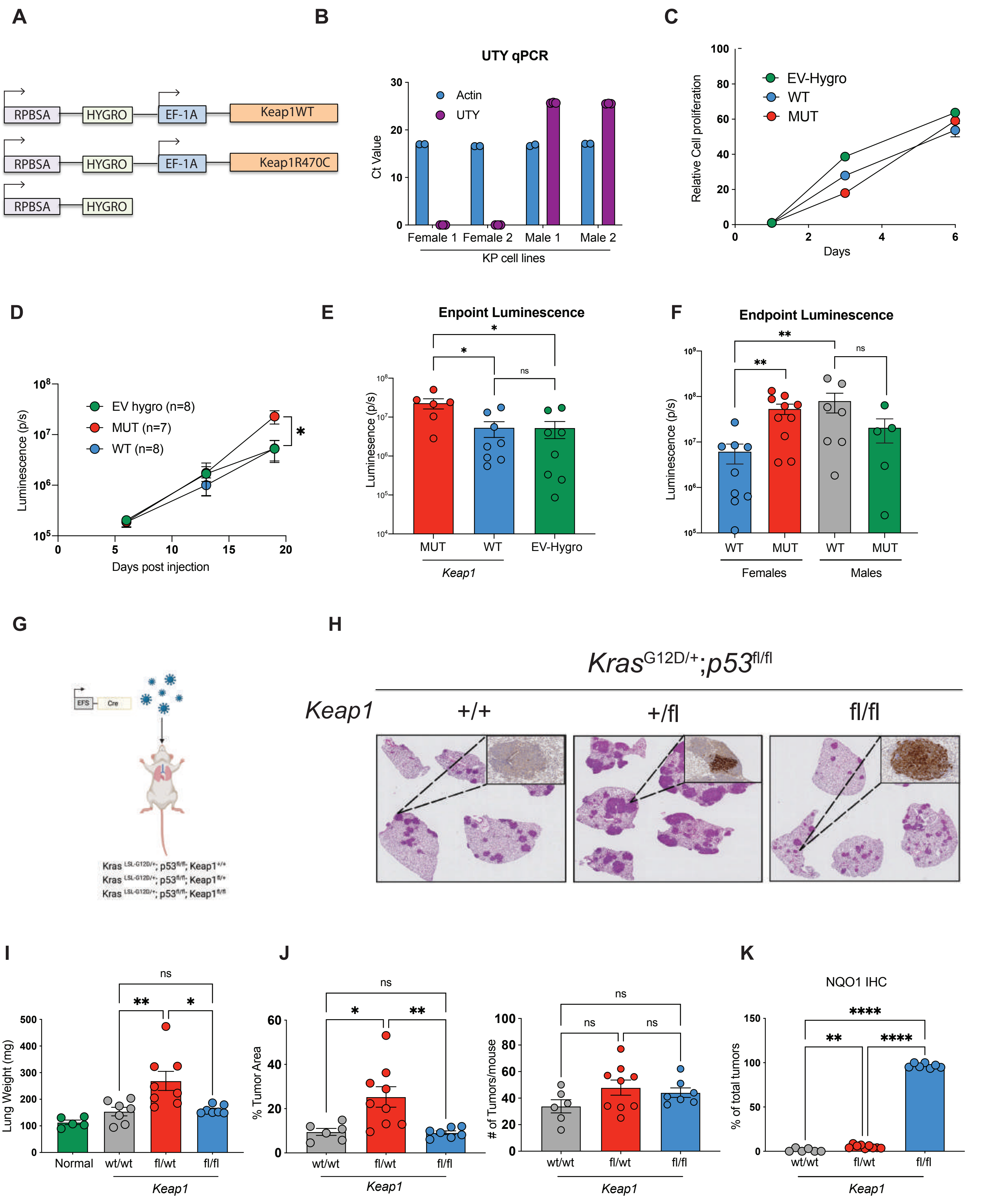
A
RPBSA
HYGRO
EF-1A
Keap1WT
RPBSA
HYGRO
EF-1A
Keap1R470C
RPBSA
HYGRO
B
UTY qPCR
Actin
UTY
Ct Value
Female 1
Female 2
Male 1
Male 2
KP cell lines
C
EV-Hygro
WT
MUT
Relative Cell proliferation
Days
D
EV hygro (n=8)
MUT (n=7)
WT (n=8)
Luminescence (p/s)
Days post injection
E
Enpoint Luminescence
Luminescence (p/s)
MUT
WT
EV-Hygro
Keap1
F
Endpoint Luminescence
Luminescence (p/s)
WT
MUT
WT
MUT
Females
Males
G
H
KrasG12D/+;p53fl/fl
Keap1
+/+
+/fl
fl/fl
I
Lung Weight (mg)
Normal
wt/wt
fl/wt
fl/fl
Keap1
J
% Tumor Area
wt/wt
fl/wt
fl/fl
Keap1
### of Tumors/mouse
wt/wt
fl/wt
fl/fl
Keap1
K
NQO1 IHC
% of total tumors
wt/wt
fl/wt
fl/fl
Keap1

Fig. S1

**Supplementary Fig 1: Impact of *Keap1* genetic inactivation on tumor growth in mouse models of LUAD.**

(A) Schematic of vectors overexpressing wild-type or mutant *Keap1* and empty vector control use to modify cell lines in the orthotopic LUAD model. (B) Gene expression of UTY in KP cell lines isolated from female or male animals. Cell line male 2 was selected for subsequent experiments (C) *In vitro* proliferation analysis of KP cell lines transduced with vectors shown in B reveals no difference in growth kinetics (D) Growth kinetics analysis of KP cells expressing *Keap1* wild-type, mutant or empty vector control In C57BL/6J female hosts reveals increased proliferation by *Keap1* mutant cells (E) Endpoint luminescence of a representative experiment outlined in D. (F) Endpoint luminescence of *Keap1* wild-type and mutant tumors established in C57BL/6J female and male hosts. (G) Schematic diagram of KP *Keap1*$^{+/+}$, *Keap1*$^{fl/+}$ or *Keap1*$^{fl/fl}$ GEMM mice infected with 20K TU *Cre*-expressing lentivirus. (H) Representative H&E and NQO1 immunohistochemical staining of Keap1$^{+/+}$, *Keap1*$^{fl/+}$ and *Keap1*$^{fl/fl}$ KP autochthonous tumors. (I) Lung weight (J) Tumor area quantification, number of tumors and (K) NQO1 staining quantification of *Keap1*$^{+/+}$, *Keap1*$^{fl/+}$ and *Kea*p1$^{fl/fl}$ KP GEMM mice infected with *Cre*-expressing lentivirus 3.5 months post infection. Each symbol represents an individual mouse. Each experimental subgroup had n≥6 mice. *P<0.05; **P<0.01; ***P<0.001; ****P < 0.0001

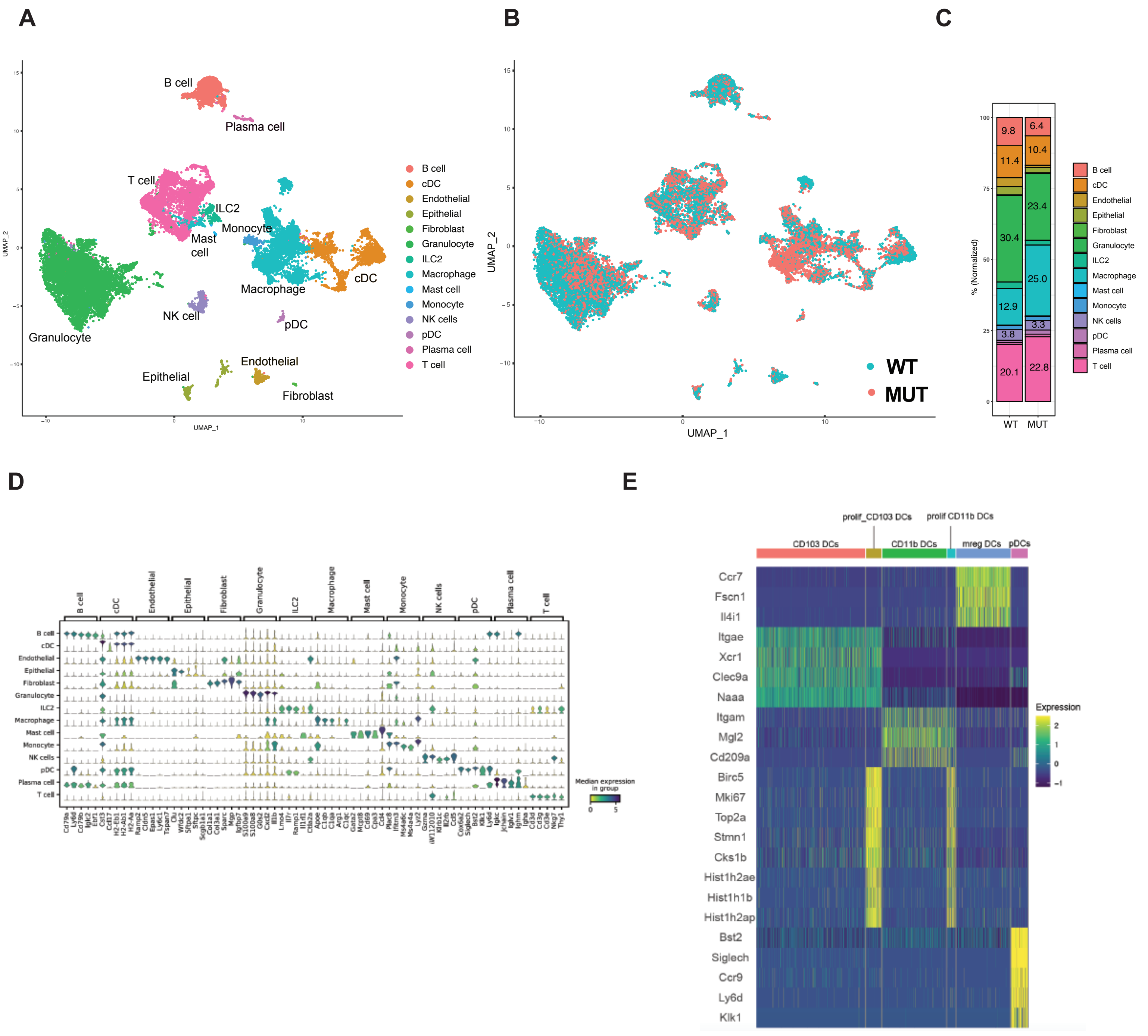
A
B
C
D
E
B cell
Plasma cell
T cell
ILC2
Monocyte
Mast cell
Macrophage
cDC
NK cell
pDC
Granulocyte
Endothelial
Epithelial
Fibroblast
UMAP_1
UMAP_2
WT
MUT
% (Normalized)
Expression
Median expression in group
CD103 DCs
prolif_CD103 DCs
CD11b DCs
prolif CD11b DCs
mreg DCs
pDCs
Ccr7
Fscn1
Il4i1
Itgae
Xcr1
Clec9a
Naaa
Itgam
Mgl2
Cd209a
Birc5
Mki67
Top2a
Stmn1
Cks1b
Hist1h2ae
Hist1h1b
Hist1h2ap
Bst2
Siglech
Ccr9
Ly6d
Klk1

Fig S2

**Supplementary Fig 2: scRNA-seq of orthotopic *Keap1* wild-type and mutant tumors.**

(A) UMAP visualization of the different immune cell lineages identified by scRNA-seq, clustered and colored by cell type. Clusters identified based on gene expression. (B) UMAP representation of the distribution of immune cell lineages in *Keap1* wild-type (blue) and mutant (red) lung tumors. (C) Bar plot showing distribution of the immune cell subsets in *Keap1* wild-type and mutant mouse lung tumors. (D) Stacked violin plots depicting the top 5 differentially expressed genes in each cluster shown in A. (E) Heatmap showing UMI counts of selected bona fide genes with key indicating cell type of origin. *$P<0.05$; **$P<0.01$; ***$P<0.001$; ****$P < 0.0001$

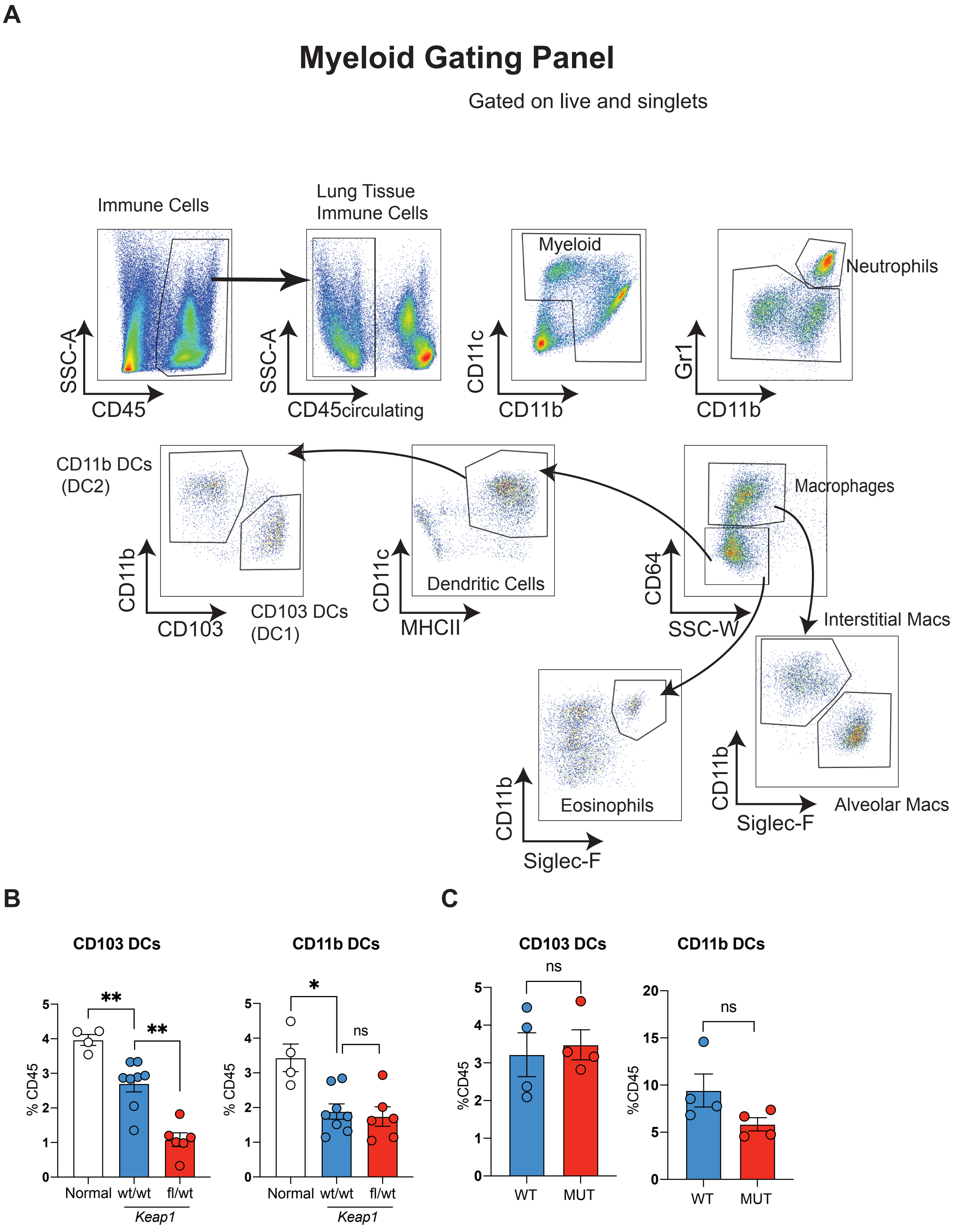
A
Myeloid Gating Panel
Gated on live and singlets
Immune Cells
Lung Tissue
Immune Cells
Myeloid
Neutrophils
SSC-A
CD45
SSC-A
CD45circulating
CD11c
CD11b
Gr1
CD11b
CD11b DCs
(DC2)
Dendritic Cells
Macrophages
CD11b
CD103
CD103 DCs
(DC1)
CD11c
MHCII
CD64
SSC-W
Interstitial Macs
Eosinophils
CD11b
Siglec-F
CD11b
Siglec-F
Alveolar Macs
B
CD103 DCs
CD11b DCs
% CD45
Normal
wt/wt
fl/wt
Keap1
**
**
*
ns
C
CD103 DCs
CD11b DCs
%CD45
WT
MUT
ns
ns

Fig. S3

**Supplementary Fig 3**: **Impact of *Keap1* loss on DCs**. (A) Gating strategy for myeloid lineage. (B) Percentage of CD103 and CD11b dendritic cells out of total tissue-infiltrating immune cells (CD45+CD45circ-) in healthy (non-tumor bearing) lungs and lungs with autochthonous *Keap1*$^{+/+}$ and *Keap1*$^{fl/+}$ tumors. Each symbol represents an individual mouse. (C) Analysis summary of orthotopic tumors showing percentage of CD103 and CD11b DCs out of total tissue-infiltrating immune cells (CD45+CD45circ-) in *Keap1* wild -type and mutant tumors established in male hosts. *P<0.05; **P<0.01; ***P<0.001; ****P < 0.0001

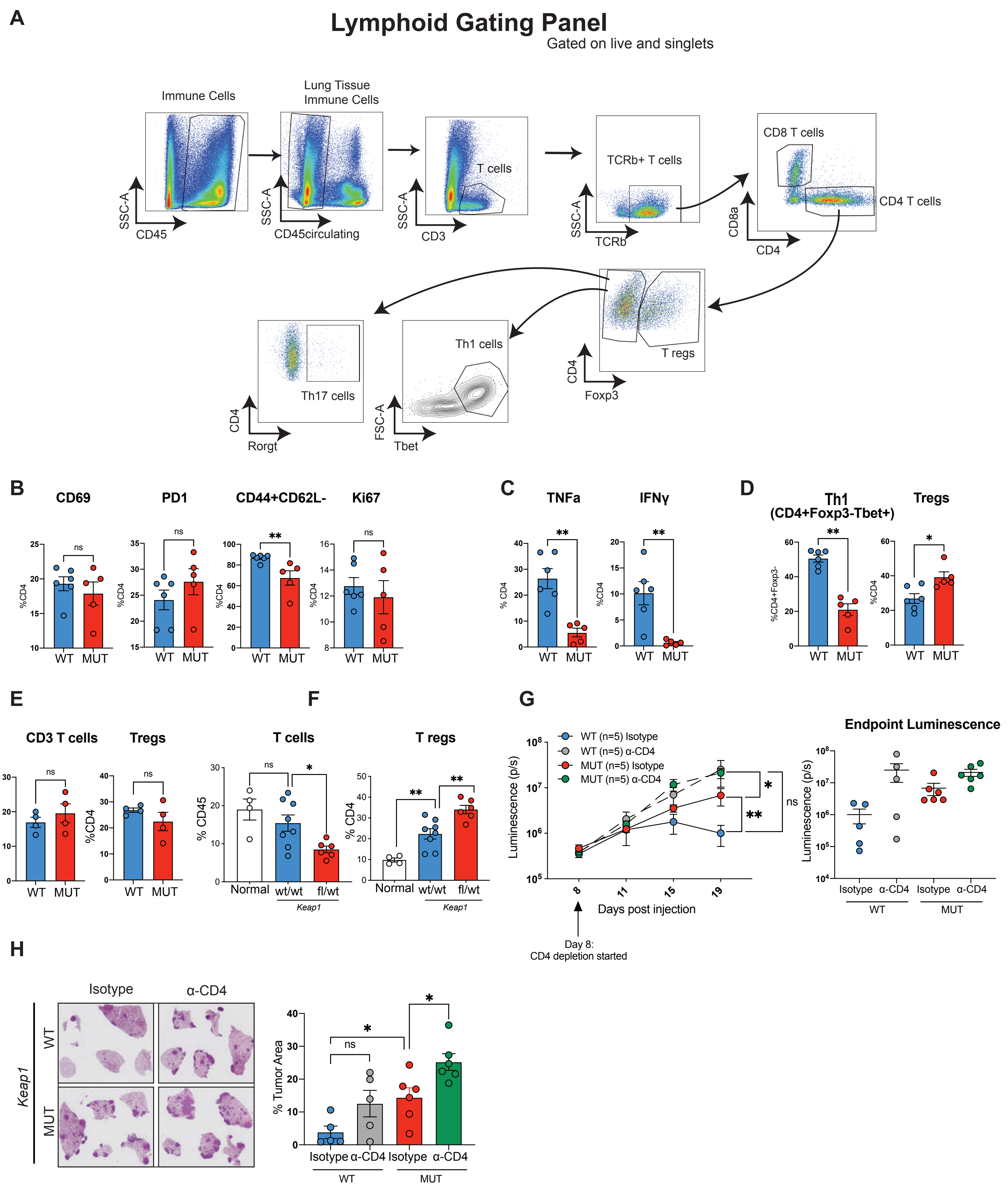
A
Lymphoid Gating Panel
Gated on live and singlets
Immune Cells
Lung Tissue Immune Cells
T cells
TCRb+ T cells
CD8 T cells
CD4 T cells
SSC-A
CD45
CD45circulating
CD3
TCRb
CD8a
CD4
T regs
Th1 cells
Th17 cells
Foxp3
Tbet
Rorgt
FSC-A
B
CD69
PD1
CD44+CD62L-
Ki67
C
TNFa
IFNγ
D
Th1 (CD4+Foxp3-Tbet+)
Tregs
WT
MUT
ns
E
CD3 T cells
F
T cells
T regs
Normal
wt/wt
fl/wt
Keap1
G
WT (n=5) Isotype
WT (n=5) α-CD4
MUT (n=5) Isotype
MUT (n=5) α-CD4
Luminescence (p/s)
Days post injection
Day 8: CD4 depletion started
Endpoint Luminescence
Isotype
α-CD4
H
Keap1
% Tumor Area

Fig. S4

**Supplementary Fig 4: *Keap1*-mutant tumors do not significantly impact CD4 anti-tumor immune responses.** (A) Gating strategy for lymphoid lineage cells. (B) Percentage of $CD69^+$, $PD1^+$, $CD44^+CD62L^-$, and $Ki67^+$ lymphocytes among CD4+ T cells in wild-type and mutant *Keap1* tumors. Each symbol represents an individual mouse. Each experimental subgroup had n≥5 mice. (C) Percentage of intracellular IFNγ and TNFα positive cells among the CD4 T lymphocytes in wild-type and *Keap1*-mutant tumors. Each symbol represents an individual mouse. Each experimental subgroup had at least 5 mice. (D) Percentage of Th1 cells (out of CD4+Foxp3-) and Tregs (out of CD4 T cells) in wild-type and *Keap1*-mutant tumors. Each symbol represents an individual mouse. (E) Percentages of T cells (out of total immune cells) and T regs (out of CD4 T cells) in *Keap1* wild-type and mutant tumors established in male hosts. (F) Percentages of T cells (out of total immune cells) and T regs (out of CD4 T cells) in autochthonous *Keap1*$^{+/+}$ and *Keap1*$^{fl/+}$ tumors. (G) Left: Growth kinetics of *Keap1* wild-type and mutant tumors in female hosts upon antibody-mediated CD4 T cell depletion. Depletion was initiated at Day 8 after verifying tumor engraftment and continued until experimental endpoint. Right: Endpoint luminescence for experiment on the left. (H) Representative images of lung tumor burden (left) and quantification (tumor area/total lung area) by H&E staining. *P<0.05; **P<0.01; ***P<0.001; ****P < 0.0001

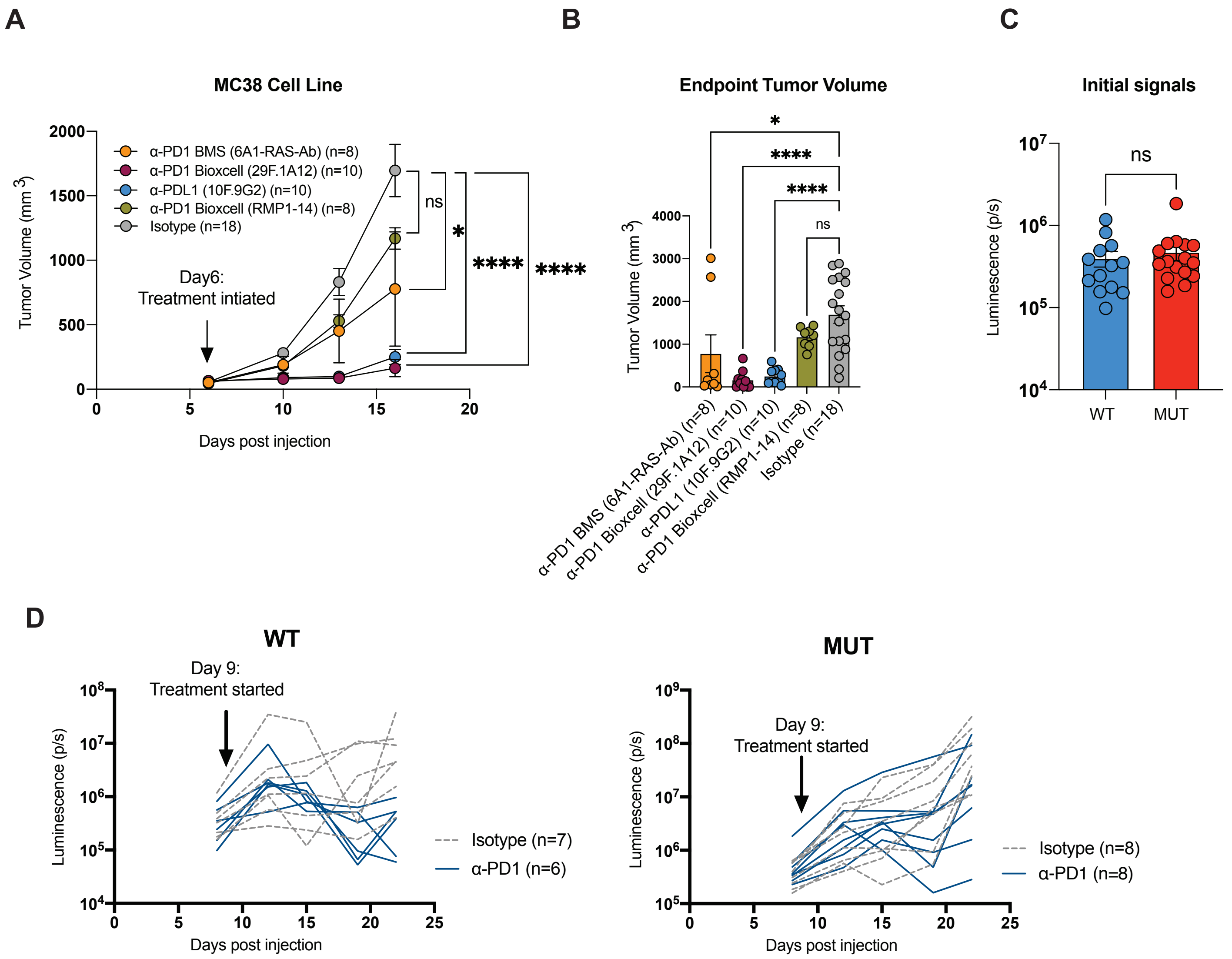
A
MC38 Cell Line
α-PD1 BMS (6A1-RAS-Ab) (n=8)
α-PD1 Bioxcell (29F.1A12) (n=10)
α-PDL1 (10F.9G2) (n=10)
α-PD1 Bioxcell (RMP1-14) (n=8)
Isotype (n=18)
Day6:
Treatment intiated
ns
*
****
****
Tumor Volume (mm 3)
Days post injection
B
Endpoint Tumor Volume
*
****
****
ns
Tumor Volume (mm 3)
α-PD1 BMS (6A1-RAS-Ab) (n=8)
α-PD1 Bioxcell (29F.1A12) (n=10)
α-PDL1 (10F.9G2) (n=10)
α-PD1 Bioxcell (RMP1-14) (n=8)
Isotype (n=18)
C
Initial signals
ns
Luminescence (p/s)
WT
MUT
D
WT
Day 9:
Treatment started
Luminescence (p/s)
Days post injection
Isotype (n=7)
α-PD1 (n=6)
MUT
Day 9:
Treatment started
Luminescence (p/s)
Days post injection
Isotype (n=8)
α-PD1 (n=8)

Fig. S5

**Supplementary Fig 5: Checkpoint blockade therapy in syngeneic mouse models.** (A) Optimization of checkpoint blockade antibodies in MC38 colon adenocarcinoma tumors. Tumor growth of MC38 cells subcutaneously injected in the right and left flanks of C57BL/6J mice was measured following treatment with various widely used immune checkpoint inhibitors. Treatment for all antibodies was initiated at Day 6, and continued until experimental endpoint, except for anti-PDL1 which was administered for a total of 3 doses. For isotype control, mice treated with various isotypes were pooled together as no differences were observed between the different isotype control antibody injections. Experimental subgroups treated with an anti-PD1 antibody or anti-PDL1 had ≥5 mice. Experimental subgroups treated with isotype control antibody had at least n≥3 mice. (B) Tumor volume at endpoint for the experiment in A. Each symbol represents an individual tumor. (C) Luminescence signals 24hrs prior to a-PD1 treatment initiation of Keap1 wild-type and mutant KP lung tumors shown in Fig. 4A. (D) Checkpoint blockade responses in individual mice for data shown in Fig. 4A.

*P<0.05; **P<0.01; ***P<0.001; ****P < 0.0001

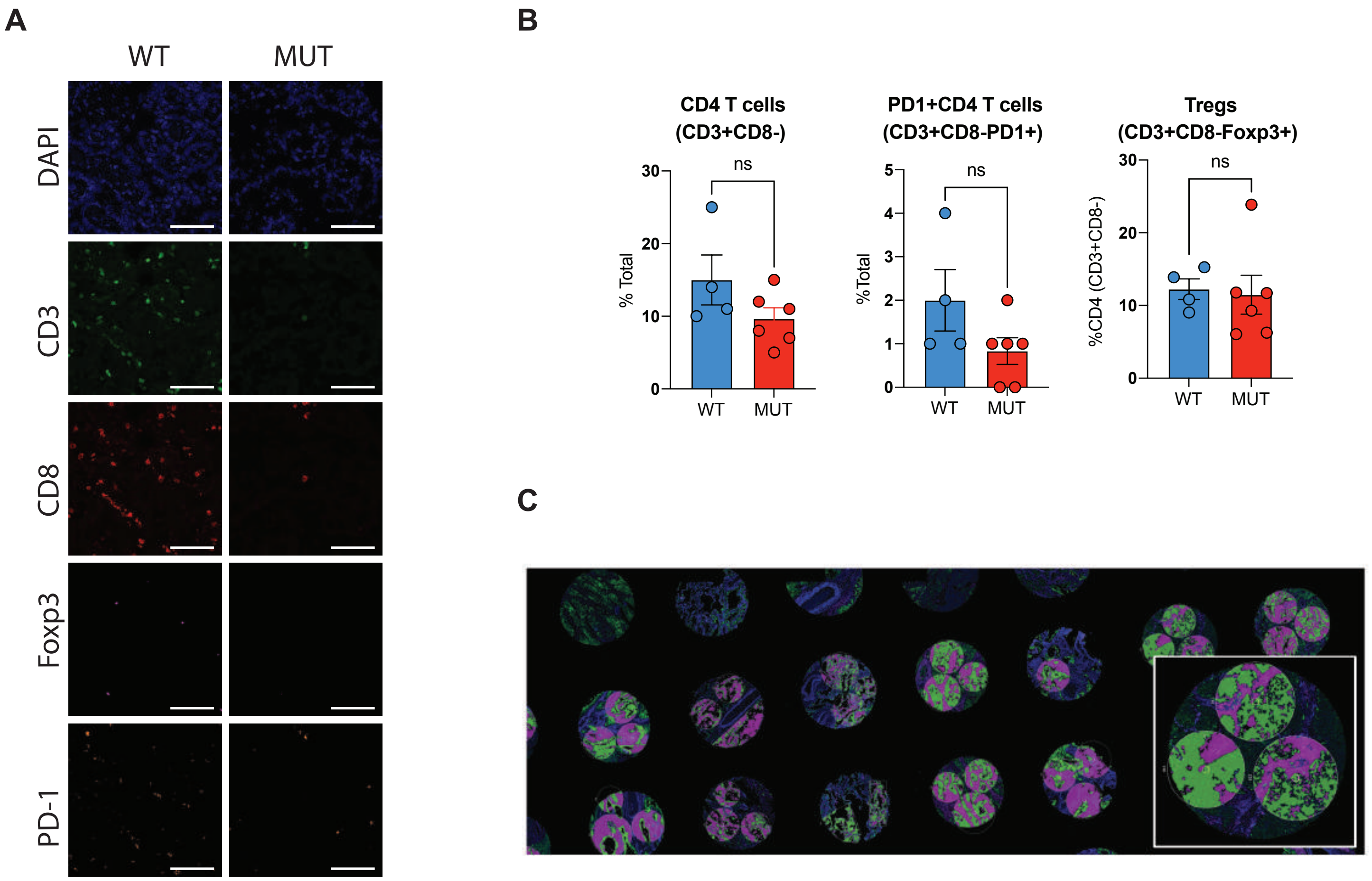
A
WT
MUT
DAPI
CD3
CD8
Foxp3
PD-1
B
CD4 T cells
(CD3+CD8-)
ns
% Total
30
20
10
0
WT
MUT
PD1+CD4 T cells
(CD3+CD8-PD1+)
ns
%Total
5
4
3
2
1
0
WT
MUT
Tregs
(CD3+CD8-Foxp3+)
ns
%CD4 (CD3+CD8-)
30
20
10
0
WT
MUT
C

Fig. S6

**Supplementary Fig 6: Multi-color immunofluorescence and Nanostring GeoMx proteomic analysis of *KEAP1* wild-type and mutant human LUAD tumors.**
(A) Representative immunofluorescence images of individual markers in *KEAP1* wild-type and mutant human tumors. Scale bar 20um. (B) Quantification of Th cells (CD3+CD8-), PD1+ Th cells (CD3+CD8-PD1+) and Tregs (CD3+CD8-Foxp3+) in *KEAP1* wild-type and mutant human tumors. Tumor area was identified based on H&E staining (C) Representative image of the LUAD tissue microarray. Three “circles” (fields of view) per tumor sample were quantified. Pancytokeratin positive staining shown in green and negative shown in magenta. *P<0.05; **P<0.01; ***P<0.001; ****P < 0.0001
